# Abolished miR158 activity leads to 21-nucleotide tertiary phasiRNA biogenesis that targets *NHX2* in *Arabidopsis thaliana*

**DOI:** 10.1101/2021.01.27.428373

**Authors:** Abhinandan Mani Tripathi, Rajneesh Singh, Akanksha Singh, Ashwani Kumar Verma, Parneeta Mishra, Shiv Narayan, Pramod Arvind Shirke, Sribash Roy

## Abstract

Small RNAs including microRNAs (miRNAs) are short 20-24-nucleotide non-coding RNAs. They are key regulators of gene expression in plants and other organisms. Some small RNAs, mostly 22-nucleotide long trigger biogenesis of secondary small interfering RNAs (siRNAs). Those siRNAs having distinctive phased configuration are known as phased siRNAs (phasiRNAs) and act either in *cis* or *trans* enhancing silencing cascade. Here, we report natural variants of *MIR158* having deletions or insertions led to negligible or reduced expression of miR158. The deletion/insertion events affected processing of primary transcript of miR158 to precursor and to mature miR158. We show that miR158 targets a pseudo-pentatricopeptide gene and its abolished activity led to 21-nucleotide tertiary phasiRNA generation from its target. The biogenesis of these phasiRNAS is triggered by *TAS2* derived two siRNAs. Accordingly, small RNA analyses of these natural variants, mutants and over-expression lines of *MIR158* or its target exhibited enhanced or reduced phasiRNA biogenesis. Finally, we functionally validated the highest expressed tertiary phasiRNA that targets *NHX2* thereby regulating transpiration and stomatal conductance. Overall, we deciphered a new module of small RNA network, miRNA-*TAS*-siRNA-pseudogene-tertiary phasiRNA-*NHX2*, suggesting an additional layer of gene regulation and larger role of pseudogene in plants.

## INTRODUCTION

Small RNAs (sRNAs) are typically 20 to 24 nucleotides long non-coding RNAs that act as important gene and chromatin regulators in plants. miRNAs are a class of small RNAs those negatively regulate gene expression either by homology mediated cleavage of mRNA or by translational inhibition affecting many aspects of plant development (Bartel, 2004). They originate from miRNA genes (*MIRNA*) transcribed by RNA polymerase II. The primary transcripts (pri-miRNA) have characteristic stem-loop like secondary structures of various lengths and complementary regions (Hirsch et al., 2006; Xie et al., 2005). The appropriate secondary structures are then processed by RNase III enzyme Dicer-like1 (DCL1) (Bologna et al., 2009; Zhang et al., 2017) forming pri-miRNA to precursor miRNA (pre-miRNA) and finally to mature miRNA duplex (Voinnet, 2009). One strand of the mature miRNA duplex gets incorporated into ARGONAUTE (AGO) protein to form a RNA-induced silencing complex (RISC) and carry out either post-transcriptional gene silencing or translational inhibition (Bartel, 2004).

Besides target degradation or translational inhibition, some miRNAs also act as a trigger for biogenesis of another class of small RNAs, the tasiRNAs (*trans* acting small interfering RNAs) or phasiRNAs (phased small interfering RNAs) to target downstream genes (Allen et al., 2005; Yoshikawa et al., 2005). The loci that generate such siRNAs are known as *PHAS* loci (Liu et al., 2020). The triggering at the target *PHAS* locus can be mediated either by a single 22-nt miRNA (one hit model) or by two 21-nt miRNAs (two hit model) (Axtell et al., 2006; Fei et al., 2013). After cleavage, the 3’ fragment is converted to dsRNA by RNA DEPENDENT RNA POLYMERASE6 (RDR6) with the assistance of AGO1-RISC or AGO7-RISC and SUPPRESSOR OF GENE SILENCING3 (SGS3). This dsRNA is then sequentially processed by DCLs with the help of DOUBLE-STRANDED RNA BINDING FACTORS into 21-or 24-nucleotides secondary siRNAs from the 5’ end of the template strand (Adenot et al., 2006; Vazquez et al., 2004). In two hit model, well studied in *Arabidopsis TAS3*, although two miR390 bind to the *PHAS* locus, but only the 3’ proximal end is cleaved while 5’ binding site defines the boundary making tasiRNA production efficient, precise and accurate (Axtell et al., 2006; Liu et al., 2020). However, in gymnosperms and eudicots the 5’ miR390 target sites are cleavable likely generating tasiRNAs in 5’ to 3’ direction (de Felippes et al., 2017). The bidirectional processing mechanism has not been reported in *Arabidopsis* (Xia et al., 2017). However, using artificial *trans*-acting siRNA transcript constructs the twin cleavage was reported in *Arabidopsis* but with poor phasing of siRNAs (de Felippes et al., 2017).

*PHAS* loci can be located both within protein-coding and non-coding genes (Fei et al., 2013). Among the non-coding *PHAS* loci, the *TAS* loci, their targets and derived phasiRNA have been well studied. In model plant *Arabidopsis thaliana (A. thaliana), TAS1a, TAS1b, TAS1c* and *TAS2* are targeted by miR173, and their 3’ cleavage products produce either identical or very closely related tasiRNAs (Yoshikawa et al., 2005). *TAS3* and *TAS4* are targeted by miR390 and miR828, respectively and derived tasiRNA target three auxin response factors: ARF2, ARF3, and ARF4 (Allen et al., 2005; Williams et al., 2005) and MYB transcription factors (Xia et al., 2012). Non-coding *PHAS* loci have been reported from other plant species as well (Huang et al., 2019; Xia et al., 2015). On the other hand, *PHAS* loci within protein-coding genes are leucine-rich repeat (*NLR*) genes (Zhai et al., 2011), *MYB* transcription factor (TF) genes (Xia et al., 2013), *AUXIN RESPONSE FACTOR* (*ARF*) genes (Xia et al., 2017), *NAC* TF genes (Sosa-Valencia et al., 2017) and *PENTATRICOPEPTIDE REPEAT (PPR)* genes (Chen et al., 2007; Howell et al., 2007) etc. Importantly, some *PPR* also produces tertiary siRNAs whose production is triggered by tasiRNAs derived from *TAS* or *TAS-*like genes (Howell et al., 2007; Xia et al., 2013). This pathway of miRNA-*TASL-PPR*-siRNA was reported to be conserved in many plant species (Xia et al., 2013). Thus it is still unexplored if miRNA acts as a negative regulator of phasiRNA biogenesis. Moreover, though the biological functions of different phasiRNAs are well documented (Marin et al., 2010; Shahid et al., 2018; Zhou et al., 2013), the role of tertiary phasiRNAs is yet to be illuminated (Jia et al., 2020).

miR158 is amongst the first released 16 plant miRNAs (Reinhart et al., 2002). It is a non-conserved young miRNA, found only in members of Brassicaceae (Supplemental Figure 1A) (Fahlgren et al., 2007; Rajagopalan et al., 2006). Later it was identified in several cruciferous plants such as *Arabidopsis lyrata* (Fahlgren et al., 2010), *Brassica oleracea* (Wang et al., 2012), *Raphanus sativus* (Wang et al., 2015) and *Brassica napus* (Xu et al., 2012). In *A. thaliana*, it has two family member *ath-miR158a* and *ath-miR158b* (Bartel, 2004) (Supplemental Figure 1B). More recently it has been shown that miR158 play role in pollen development by regulating PPR in *Brassica campestris* (Ma et al., 2017).

*A. thaliana* harbors wide genetic variations among wild types and have been explored extensively over the decades (Alonso-Blanco and Koornneef, 2000; Mitchell-Olds and Schmitt, 2006). Effects of single nucleotide polymorphisms in the *MIRNA* have also been reported (Ryan et al., 2010; Todesco et al., 2012). In our previous study, we observed negligible expression of miR158 in a particular *A. thaliana* population of Indian west Himalayas under different growing conditions (Tripathi et al., 2019). These populations are reported from elevation range of 700 m above mean sea level (amsl) to 3400 m amsl having varied climatic conditions. Here, we show that negligible or moderate expression of miR158 is due to deletion and insertion events respectively, in *MIR158* affecting it’s processing to mature miRNA. miR158 targets a pseudo-*PPR*. However, null expression of miR158 led to enhanced tertiary phasiRNAs biogenesis from pseudo-*PPR* via *TAS2* derived two siRNA triggers. Finally, we show that one of the tertiary phasiRNAs targets *NHX2* thereby regulating physiological processes prompting plants to complete life cycle early, presumably as a strategy of local adaptation.

## RESULTS

### Population specific expression of miR158

Our previous analyses of small RNA profiling of the three populations Deh, Mun, and Chit showed negligible expression of miR158 in Mun as compared to the other two populations, both under field and controlled growth conditions (Table S4 of(Tripathi et al., 2019)). To reconfirm the negligible expression of miR158 in Mun, we analyzed its expression under native, controlled and common garden conditions along with miR158 SALK mutant using quantitative real time PCR (qRT-PCR). The expression of miR158 was consistently negligible in Mun under controlled, native and common garden conditions and as expected, in miR158 mutant (Figure1A; Supplemental Figure 2A and 2B) as compared to the other two populations. We also checked the relative expression pattern of miR158a, miR158b, and miR158a-3p arm and miR158a-5p arm under controlled condition. Data suggested the relative expression of miR158a and miR158-3p was significantly dominant over miR158b and miR158-5p respectively, in all the populations (Supplemental Dataset 1; Supplemental Figure2C and 2D).

### Polymorphisms in *MIRNA158* are unique to the west Himalayan populations

To find out probable reason of population specific expression of miR158, we first analyzed the putative promoter sequence of miR158 (one kb upstream of transcription start site, TSS) and observed no significant sequence polymorphisms amongst the three populations, ruling out the low expression of miR158 in Mun was due to differential promoter activity (Supplemental Figure 3). However, PCR amplification and sequencing of *MIRNA158 (MIRNA158a)* from a few plants of each of the three populations exhibited length polymorphisms (Figure 1B and 1C). Subsequently, ∼50 plants from each of the three populations were analyzed (Supplemental Figure 4). The results indicated that while the 535 base pair (bp) *MIRNA158* of reference Col-0 (Szarzynska et al., 2009) (hereafter NTL, Normal Type Locus) was present in all Chit plants, the Mun population harbors two different lengths of *MIRNA158*, a 437 bp locus having discontinuous deletion of total 98 bp (starting at 10 bp downstream of transcription start site, TSS including four nucleotides of 5’ precursor; hereafter DTL, Deleted Type Locus) and a760 bp locus with an insertion of 225 bp (10 bp downstream of TSS; here after ITL, Inserted Type Locus). Notably, the frequency of occurrence of DTL in Mun was 0.94, remaining being ITL, and Deh harbors both NTL and ITL having frequency of 0.5 each (Figure 1B and 1D; Supplemental Figure 5).

**Figure 1:**
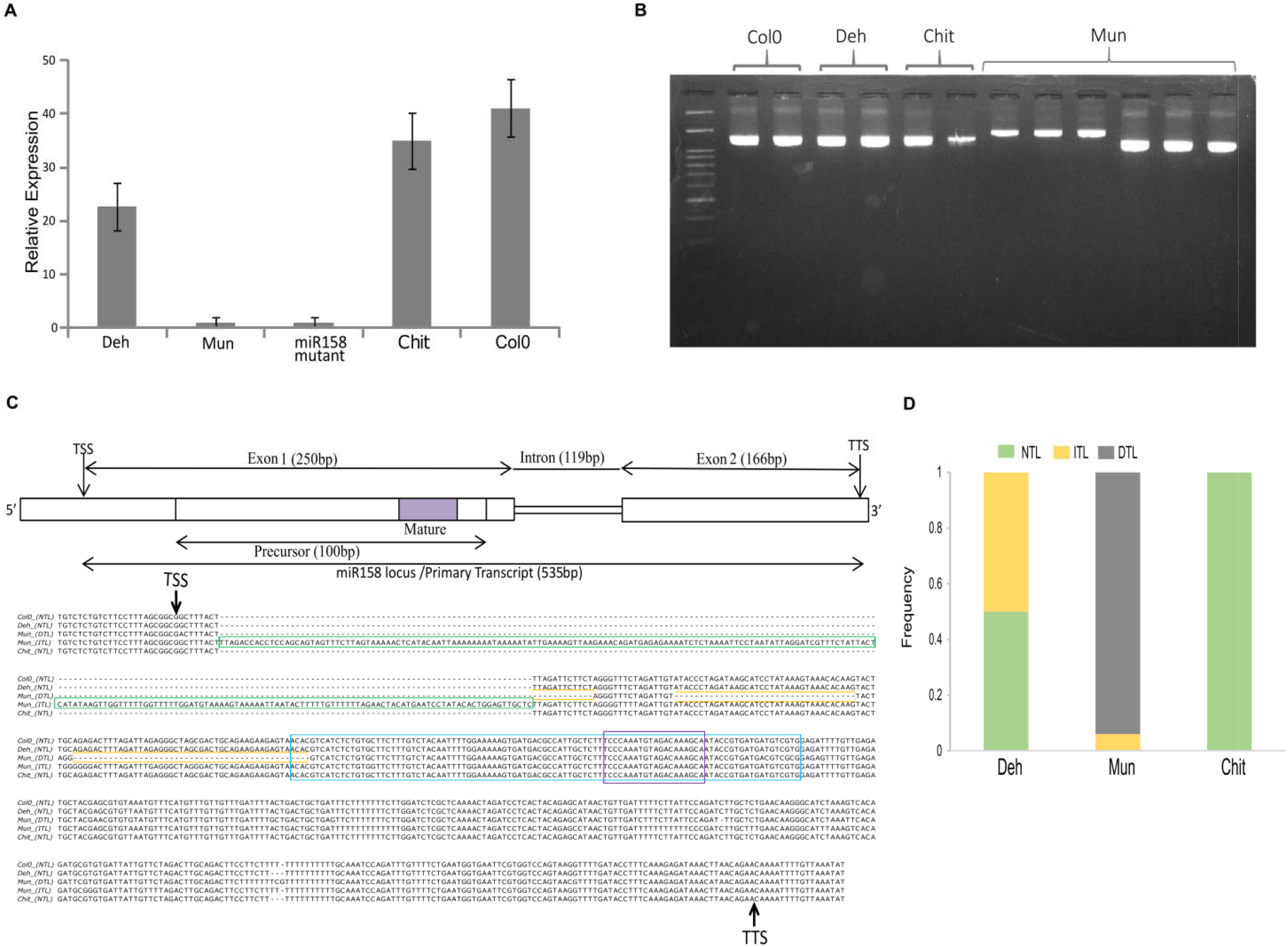
Expression pattern and length polymorphisms of miR158 locus in different populations. **(A)** The expression pattern of miR158 in different west Himalayan populations as well as miR158 SALK mutant and Col-0 were analyzed by qRT-PCR. The data represents the mean of four independent biological replicates ± SD. **(B)** Length polymorphism of miR158 locus as revealed by agarose gel electrophoresis of the PCR amplification products from a few individuals of each of the three populations. While Deh, Chit and Col-0 exhibited same fragment length as expected, the Mun exhibited two types of fragment lengths due to insertion and deletion. **(C)** Multiple alignment of miR158 locus sequenced from the three populations. The insertion sequence in ITL, insertion type locus and the deleted sequence in DTL, deleted type locus are shown in green and yellow box, respectively. The precursor and mature sequences of miR158 are shown in blue and violet boxes, respectively. TSS, transcription start site and TTS, transcription termination site are indicated by arrows. The locus was annotated against Col-0 reference locus and schematically represented above the aligned sequences. **(D)** Frequency of insertion/deletion in miR158 locus in west Himalayan *Arabidopsi* populations as analyzed from 50 plants of from each population.

To examine if the insertion/deletion polymorphisms were present in the primary transcript as well, we PCR amplified the locus using cDNA as template. Amplified product exhibited similar length polymorphisms as observed with genomic DNA, ruling out the observed polymorphisms was due to any splice variants (Supplemental Figure 6). We checked if these polymorphisms are unique to the west Himalayan populations by comparing the *MIRNA158* sequences from available 1001 genomes of *A. thaliana* (Alonso-Blanco et al., 2016). The observed insertion/deletion polymorphisms are unique to the west Himalayan populations. Further, the inserted fragment was not a transposable element (TE), as BLAST with TE database (TAIR) did not retrieve any TE. However, the inserted fragment showed 97% similarity with a sequence of chromosome four while *MIRNA158* is located on chromosome three of *A. thaliana*. Thus, the insertion event may be due to inter-chromosomal translocation.

### Polymorphisms in *MIRNA158* affect its expression pattern

We checked the expression pattern of miR158 in NTL, ITL and DTL by qRT-PCR. Data suggested the expression of miR158 was the highest, intermediate, and negligible in NTL, ITL and DTL, respectively (Figure 2A; Supplemental Dataset 2). In addition, we determined the expression level of both pri- and pre-miR158. As expected, the level of pri-miR158 was more or less similar in all the three variants (Figure 2B). However, pre-miR158 level was high, intermediate, and low in NTL, ITL and DTL, respectively as observed with mature miR158 (Figure 2C). It further suggests that the variations in the expression of miR158 was not due to promoter activity but presumably, may be due to differential processing of primary transcripts to precursors. We evaluated the secondary structures of all the three different primary transcript sequences from NTL, ITL and DTL plants. The secondary structure prediction of all the variants revealed that the lower stem and their internal loops in DTL plants were structurally different (disturbed) from the canonical ones and so was the large internal loop in ITL plants but these were intact in NTL plants (Figure 2D - 2F). However, there were no such structural anomalies in the predicted secondary structures of precursor sequences of all the three variants (SupplementalFigure7). Any anomaly in the lower stem and internal loop structure can lead to sloppy primary transcript processing (Mateos et al., 2010; Zhu et al., 2013).

**Fig. 2:**
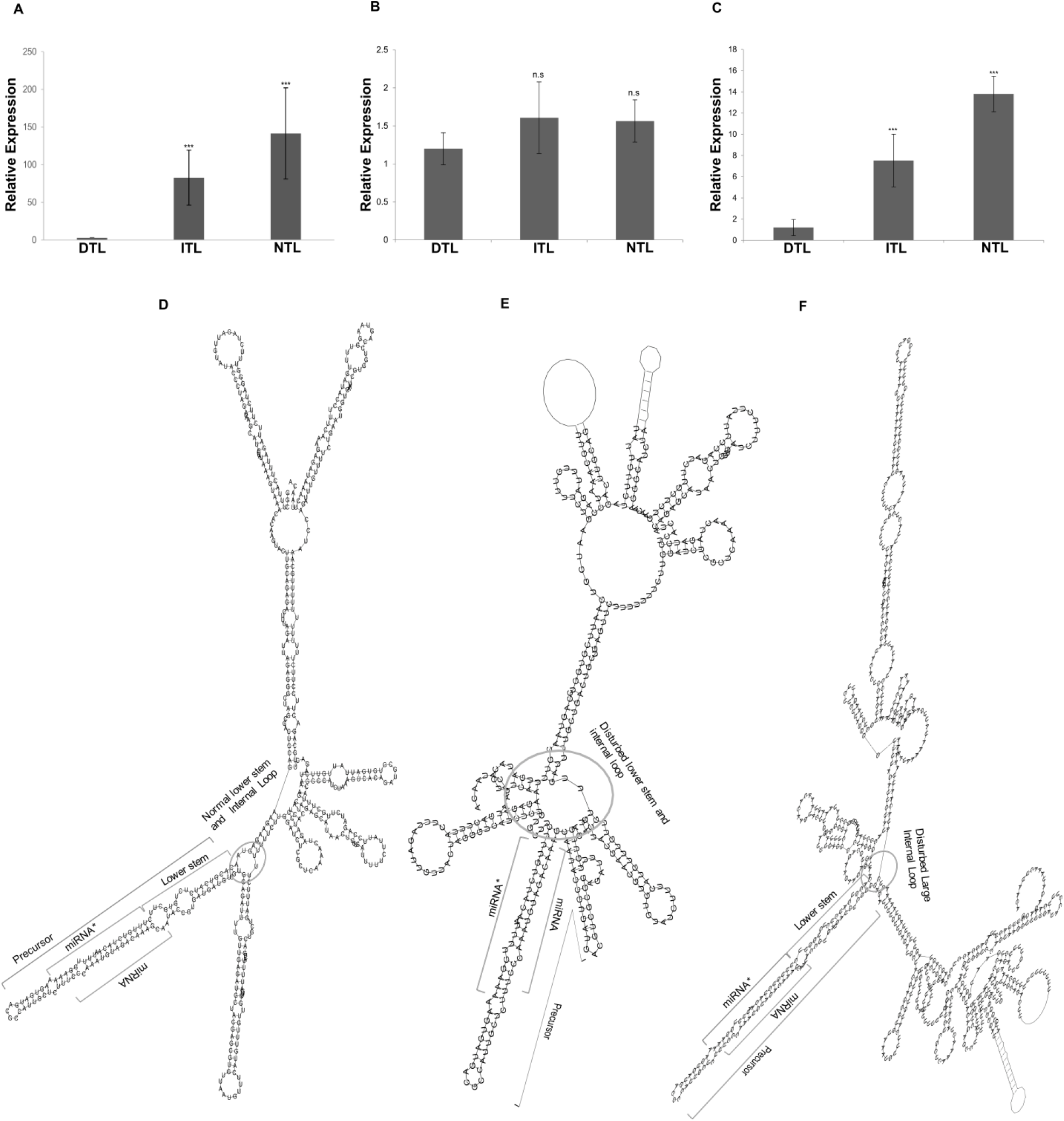
Expression patterns of miR158, pri-miR158, pre-miR158 and predicted secondary structures of pri-miR158. Relative expression pattern of: **(A)** miR158; **(B)** pri-miR158; and **(C)** pre-miR158, in DTL, ITL and NTL plants. The data represents the mean of three independent biological replicates ± SD. (***P<0.001, ** P ≤ 0.01. * P ≤ 0.05, (Student’s t-test); ns, non-significant). The predicted secondary structures of pri-miR158 in (**D)** NTL; **(E)** DTL; and **(F)** ITL, derived from the respective sequence using RNAfold web server. The miRNA/miRNA*, precursor, lower stem sequences are shown along with disturbed large internal loops in DTL and ITL (encircled).

### miR158 targets a pseudogene, pseudo-*PPR*

The targets of miR158 were predicted using psRNATarget (Dai and Zhao, 2011). Four genes namely, pseudo-pentatricopeptide repeat (pseudo*-PPR*), tetratricopeptide repeat (*TPR*), transducing/WD40 repeat (*BUB3*.*2*) and pentatricopeptide repeat (*PPR*) proteins were identified as the putative targets of miR158 (Supplemental Dataset 3). We attempted to validate the predicted targets by modified 5’ RLM-RACE and it identified pseudo-*PPR* (AT1G62860.1) as the only target of miR158 (20/20 clones). Quantification of the pseudo-*PPR* transcripts by qRT-PCR also showed inverse expression pattern with miR158 (Figure 3A and 3B), thus confirming it as the target of miR158. Further, the available degradome data (GSE55151) (Thatcher et al., 2015) analyses (P value < 0.0001) also retrieved the identified pseudo-*PPR* as the target of miR158 (Supplemental Figure 8). Our repeated attempts failed to identify any of the other predicted genes as the true target of miR158.

**Fig. 3:**
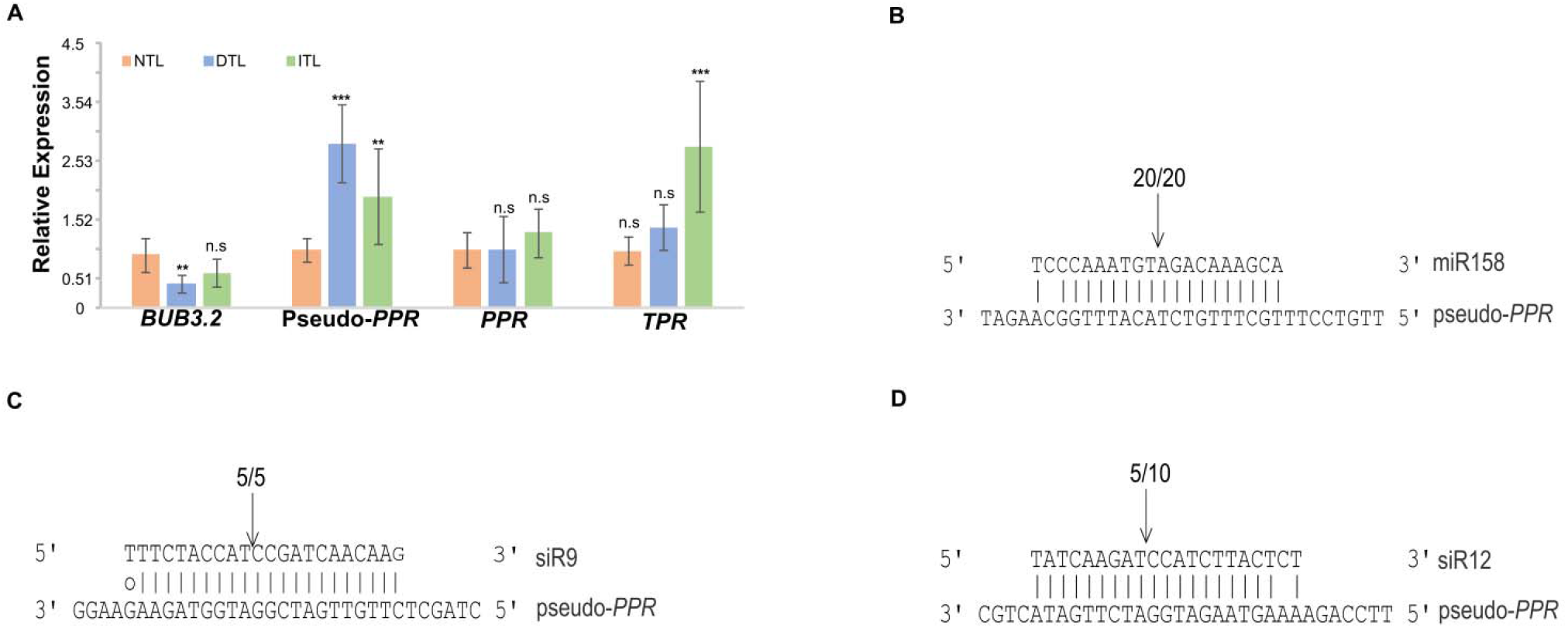
miR158 target expression pattern and target validation of miR158 and tasiRNA. **(A)** The expression pattern of all the predicted targets of miR158 as evaluated by qRT-PCR. The data represents the mean of three independent biological replicates ± SD (***P<0.001, ** P ≤ 0.01. * P ≤ 0.05, (Student’s t-test), ns (not significant)). Pseudo-*PPR* as the target of **(B)** miR158; **(C)** siR9; and **(D)** siR12, validated using modified 5’ RLM-RACE. Pseudo-*PPR*mRNA: miRNA/tasiRNA alignments are shown along with the detected cleavage site (shown by arrow). The number above arrow indicates the positive clones at this cleavage site/total number of colonies sequenced.

### The target pseudo-*PPR* acts as a PHAS locus in absence of miR158 activity

Pseudogenes have been reported to act as a sponge for some miRNAs and thus protect the cognate target from degradation (Yu et al., 2014). However, our observation that miR158 cleaves the pseudogene prompted us for detail investigation. The target pseudo-*PPR* besides harboring several stop codons also contains an open reading frame of 280 amino acid residues (Supplemental Figure 9). The parental *PPR* (AT1G64100.1) shares 78% identity with the target pseudo-*PPR*. The PPRs are mostly known to involve in RNA editing (Yagi et al., 2013) and some act as PHAS loci (Howell et al., 2007). PPRs are separated into two major classes based on the nature of their PPR motifs, the P and PLS classes (Schmitz-Linneweber and Small, 2008; Xia et al., 2013).Phylogenetic analysis of the target pseudo-*PPR* with 297 *A. thaliana PPR*s (Lurin et al., 2004) suggested it is a P-type *PPR* (Supplemental Figure 10). Being a pseudogene, we ruled out the possibility of its involvement in RNA editing and attempted to find if it acts as a PHAS locus. PHAS loci are characterized by being targeted either by one 22-nucleotide small RNA (one hit model) or by two 21-nucleotide small RNAs (two hit model) (Fei et al., 2013). We analyzed the available degradome data to determine the putative small RNAs that might cleave the pseudo-*PPR*. Our analysis indicated that apart from miR158, the pseudo-*PPR* is also the putative target of tasiRNAs, atTAS2 (2)-siR9(-) (hereafter after referred as siR9) and atTAS2(2)-siR12(-) (hereafter after referred siR12) derived from *TAS2* (Zhang et al., 2014) (Supplemental Figure 8). We validated the target cleavage by both the tasiRNAs using modified 5’ RLM-RACE and detected both the cleaved ends (Figure 3C and 3D).

Next, we mapped our small RNA libraries against the pseudo-*PPR*. A large number of pseudo-*PPR* mapped small RNAs (90%) lied between siR9 and siR12 binding sites (Figure 4A). Moreover, the small RNAs mapped between siR9 and siR12 were in phase with phasi score of 1.38 to 35.0 (P value < 0.0001) (Figure 4B) starting with the cleavage site of siR9 (702 nucleotides, from 5’ end of pseudo-*PPR)* (Figure 4C). Although siR9 is reported to be 21-nucleotide (Howell et al., 2007; Zhang et al., 2014; Zhong et al., 2013) but our small RNA analysis suggested higher expression of 22-nucleotide siR9 than the 21-nucleotide ones and siR12 were of 21-nucleotide (Figure 4E and Supplemental Table 1). Further, when small RNAs of each of the populations were mapped individually against the pseudo-*PPR*, higher number of small RNAs mapped between siR9 and siR12 binding sites in Mun than the other two populations (Supplemental Figure 11A). Additionally, out of 3138, a total of 830 non-redundant phasiRNAs from all the populations were identified those were derived from the pseudo-*PPR* (here after referred as PDP) (Supplemental Dataset 4). Majority of the PDPs were 21-nucleotide while a very small fraction was of 22-nucleotide (Supplemental Figure11B). Further, around 45% of these PDPs had characteristic 5’ terminal uridine (Supplemental Figure 11B). Since many *PPR*s are known to act as PHAS loci and phasiRNAs produced from them may show redundancy due to repetitive domains, we speculated that the PDPs may be identical with the *PPR* derived phasiRNAs. However, when we compared our pseudo-*PPR* derived small RNAs with the small RNAs derived from each of the known 23 *PPR* PHAS loci (Howell et al., 2007), only 8% of the PDPs showed similarity with other *PPR* derived small RNAs (Supplemental Dataset 5) reconfirming pseudo-*PPR* acts as PHAS locus.

**Fig. 4:**
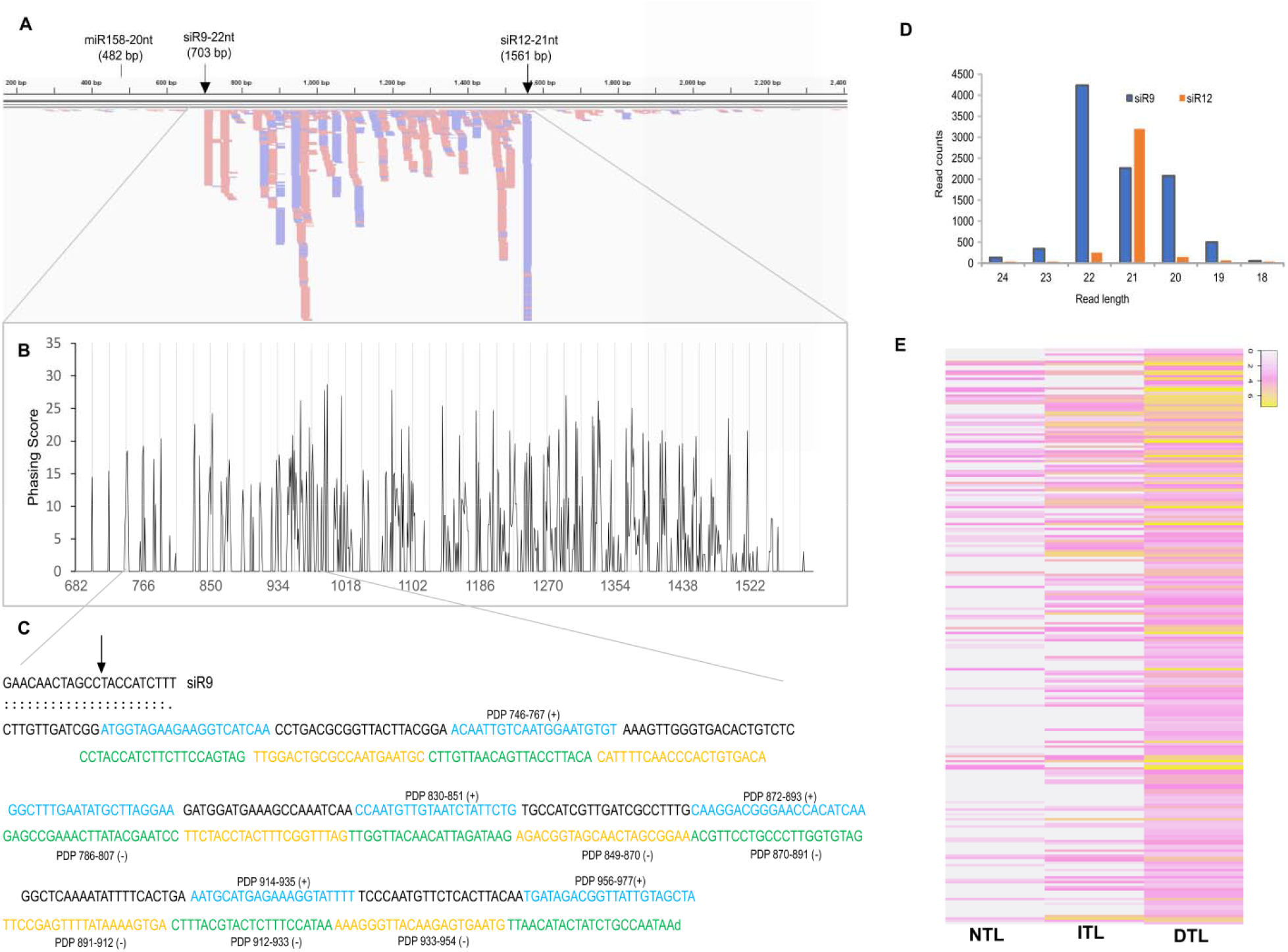
PhasiRNA analysis and expression pattern. **(A)** The small RNAs were mapped against pseudo-*PPR* shown as horizontal line along with nucleotide positions and cleavage sites (arrow) of miR158, siR9 and siR12. The red blocks depict reads aligned in forward orientation and blue in reverse orientation (visualized via Integrative Genomics Viewer (IGV)). **(B)** The phasing scores calculated using small RNAs mapped between 600-1600 nucleotide positions of pseudo-*PPR*. The X-axis and Y-axis represent the phasing score and nucleotide positions of pseudo-*PPR*, respectively. The 21 nucleotides phasing intervals are shown by vertical grey lines. **(C)** The phasing pattern of a significant cluster of phasiRNA derived from pseudo-*PPR* (PDP) initiated at siR9 cleavage site (marked by arrow). Blue and red sequences indicate the phasiRNAs (PDPs) derived from sense (+) strand while yellow and green sequence represent phasiRNAs derived from antisense (-) strand. **(D)** Read length distribution of siR9 and siR12 in Col-0. The data indicates the expression of 22-nucleotide is dominating for siR9. **(E)** Heatmap depicting expression pattern of phasiRNAs in NTL, ITL and DTL (n=238). The log2 value of PDPs read counts are represented by the colour intensity scale. (NTL-Normal Type Locus; ITL-Inserted Type Locus; DTL-Deleted Type Locus).

It was clearly evident that the abundance of PDPs was more in Mun than Deh and Chit (P value = 0.005645, F = 4.32423) (Table 1; Supplemental Figure 12A). Since above small RNA libraries were prepared from pooled leaf samples of several plants of each population (Tripathi et al., 2019), we re-examined the data using small RNA sequence data from randomly selected individual plants of each variant types. Small RNA data from individual DTL, ITL and NTL plant reconfirmed the above observations (Table 1; Figure 4D). Similarly, PDP analyses of miR158 mutant, miR158 target pseudo-*PPR* mutant and Col-0 confirmed the higher expression of PDPs in miR158 mutants and lower in pseudo-*PPR* mutant (Table 1; Supplemental Figure 12B). Further over-expression lines of both miR158 (miR158 OE) and pseudo-*PPR* (pseudo-*PPR* OE) exhibited lower and higher expression of PDP, respectively as compared to Col-0 (Table 1; Supplemental Figure 12C). These analyses suggest miR158 negatively regulate the PDP biogenesis.

**Table 1.** Comparative read counts of PDPs and related small RNAs in different populations.

|  | Deh |  | Mun |  | Chit |  | Col-0 |  |  |  |  |  |
| --- | --- | --- | --- | --- | --- | --- | --- | --- | --- | --- | --- | --- |
|  | Replicate 1 | Replicate 2 | Replicate 1 | Replicate 2 | Replicate 1 | Replicate 2 | Replicate 1 | Replicate 2 |  |  |  |  |
| miR166e-3p | 15934 | 11399 | 10459 | 10991 | 11356 | 12860 | 15400 | 7339 |  |  |  |  |
| miR158 | 60210 | 17443 | 3848 | 4707 | 23242 | 27121 | 70440 | 37725 |  |  |  |  |
| siR9 | 595 | 792 | 614 | 743 | 337 | 391 | 740 | 428 |  |  |  |  |
| siR12 | 887 | 221 | 548 | 579 | 480 | 412 | 664 | 397 |  |  |  |  |
| PDPs expression | 172 | 105 | 279 | 288 | 60 | 75 | 107 | 62 |  |  |  |  |
|  | Col-0 (NTL) |  | miR158mutant |  | Pseudo-PPRmutant |  | DTL | ITL |  | Col-0 | miR158OE | Pseudo-PPROE |
|  | Replicate 1 | Replicate 2 | Replicate 1 | Replicate 2 | Replicate 1 | Replicate 2 | Replicate 1 | Replicate 1 | Replicate 2 | Replicate 1 | Replicate 1 | Replicate 1 |
| mir166e-3p | 16026 | 17135 | 17941 | 18588 | 16318 | 17456 | 18355 | 23821 | 14525 | 54950 | 35107 | 41774 |
| miR158a-3p | 224638 | 164825 | 3208 | 2853 | 230162 | 344228 | 687 | 25228 | 42200 | 308014 | 2991449 | 389142 |
| siR9 | 1573 | 1231 | 1789 | 2231 | 1691 | 2679 | 1975 | 1083 | 2210 | 3061 | 2256 | 2466 |
| siR12 | 734 | 258 | 1225 | 833 | 470 | 1125 | 609 | 600 | 414 | 991 | 1016 | 1025 |
| PDPs expression | 174 | 184 | 241 | 265 | 113 | 129 | 552 | 419 | 386 | 228 | 169 | 1641 |

Expression (read counts) of miR158, tasiRNAs and PDPs in different west Himalayan populations and those of Col-0, SALK mutants and over expression lines along with DTL and ITL are shown in above and lower panel, respectively. The expression of miR166 was considered as technical standard being not associated with PDPs and related siRNAs pathway.

### The highly expressed PDP targets *NHX2*

Most of the 21-nucleotide siRNAs are known to involve in AGO1 mediated cleavage of target mRNA (Deng et al., 2018). In order to identify if any PDPs were associated with AGO proteins, we took advantage of previously published AGO1 and AGO4 bound small RNA data sets (GSE28591) (Wang et al., 2011). We identified a total of 200 PDPs identical with AGO1-associated and 25 PDPs were identical with AGO4-associated small RNAs (Supplemental Dataset 6). It presumably indicates a large number of the PDPs might involve in cleavage mediated functions rather than involve in RdDM pathway. For further characterization of PDPs, we focused on 11 PDPs which were derived from a significant cluster (Start: 746; End: 997; P value < 0.00001, n=11, k=236) (Supplemental Dataset 7) and were in phase with siR9 cleavage site. To find out the cleaved ends of mRNAs by these PDPs, available degradome data was analyzed. It identified three different mRNAs belonging to degradome category-2 (P value < 0.05), which indicates >1 raw read mapped at the position and abundance at position is less than the maximum but higher than the median for the transcript (Supplemental Dataset 8). These three mRNAs were putative target of one of the highly expressed PDP, PDP956-977 (+) (Supplemental Dataset 8; Supplemental Figure 13). The best target identified was *NHX2*, a Na^+^-K^+^/H^+^ antiporter protein expressed in stomatal guard cell that maintain turgor to regulate transpiration (Andrés et al., 2014; Barragán et al., 2012; Bassil et al., 2011). Other two targets were identified as DNAJ heat shock N-terminal domain-containing proteins (AT2G05250.1 and AT2G05230.1). Target validation using modified 5’ RLM-RACE also identified *NHX2* as the target of PDP956-977 (+) (Figure 5A). Subsequently, we checked the expression of *NHX2* in DTL and miR158 mutant plants by qRT-PCR. As expected the expression of *NHX2* in DTL and miR158 mutant was lower than other lines (Figure 5B; Supplemental Figure 14A). Notably, the expression of this gene in Mun was also found to be lower in our earlier global RNA sequencing data (Tyagi et al., 2016).

**Fig. 5:**
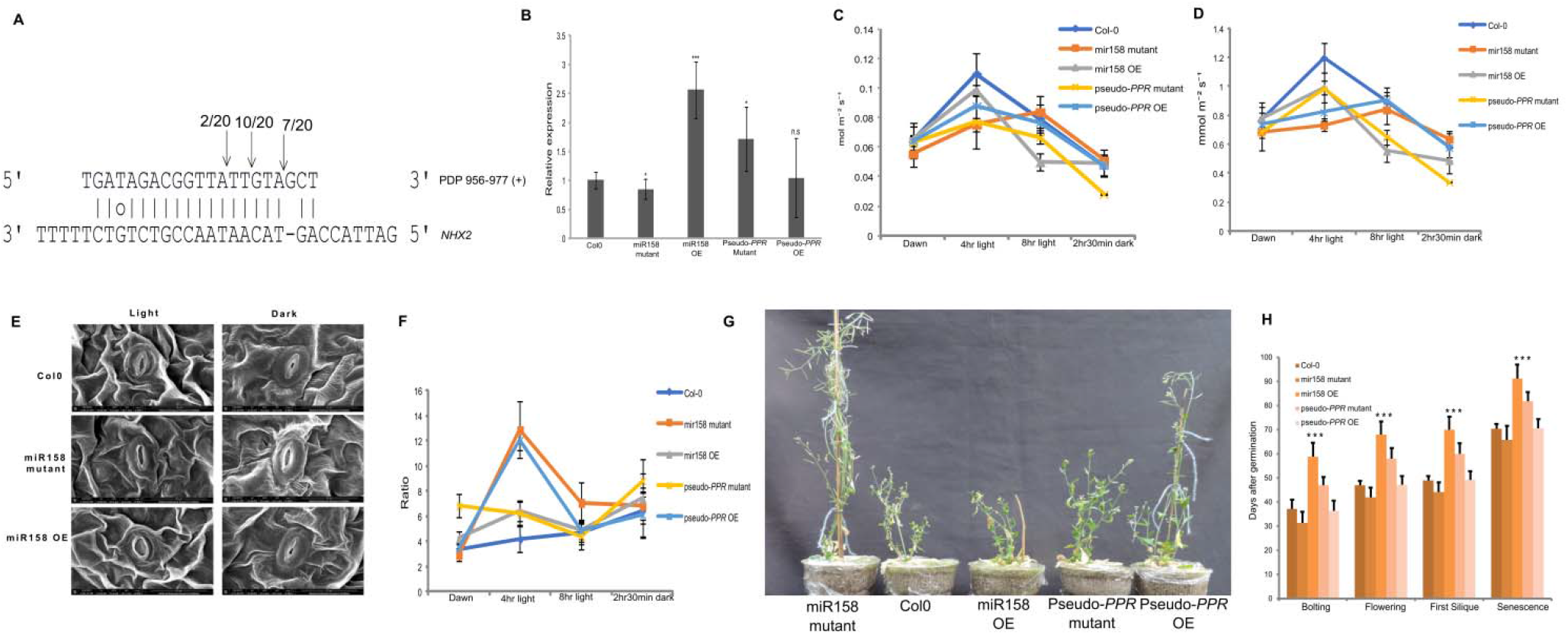
PDP target analysis and their functional characterizations. **(A)** The complementary pairing of PDP 956-977(+) with its target *NHX2* is shown with arrows indicating the cleaved positions and the numbers suggest positive clones with cleavage sites/total number of colonies sequenced. The target of the highly expressed PDP, PDP 956-977(+) validated using modified 5’ RLM-RACE. **(B)** Expression pattern of *NHX2* in different transgenics and mutant lines by qRT-PCR. The data represents the mean of three independent biological replicates ± SD. (***P<0.001,** P ≤ 0.01. * P ≤ 0.05, (Student’s t-test), ns (not significant)). Physiological parameters of mutant and transgenic lines in Col-0 background measured at different time points: **(C)** Stomatal conductance; **(D)** Transpiration rate **(E)** Scanning Electron microscope image of stomata at two time points (light and 2 hr 30 min dark) taken at 8000x **(F)** Water Use Efficiency. The data represents the mean of five biological replicated ± standard deviation. **(G)** Image of plants of miR158 mutant, miR158 OE, pseudo-*PPR* mutant, pseudo-*PPR* OE and Col0 taken on 56th day post germination. Plants were grown with normal watering till 21 days and then water was withheld for 35 days after pots were saturated with water on 21st day. **(H)** Measurement of various life cycle related parameters in mutant and transgenic lines in Col-0 background. Life cycle parameters namely, bolting, first flower opening, emergence of first silique and senescence were measured in terms of days after germination. Data represents average of 14-15 plants per genotype ± SD. Asterisk above graph shows significance of variation (‘***’ P<0.001, ‘**’ P<0.01,’*’ P<0.05, ‘.’ P<0.1, ns = not signifiant).

*NHX1* and *NHX2* are members of *NHX* gene family and are involved in regulating stomatal function (Barragán et al., 2012; Bassil et al., 2011). *NHX2* shows a diurnal expression pattern and expression of *NHX1* increases in *NHX2* mutant to counter balance the NHX2 activity (Andrés et al., 2014).We quantified the expression of *NHX1* and *NHX2* at different time points of the day (Supplemental Figure 14B and 14C). In contrast to earlier observation, we did not observe increased expression of *NHX1* in miR158 mutants, as expected with *NHX2* mutant. This may be due to knock-out effect of *NHX2* as compared to knock-down effect in miR158 mutant. Further, the maximum expression of *NHX2* was observed at 4 hr of light in Col-0 and miR158 OE while in miR158 its expression was the maximum at 8 hr of light (Supplemental Figure 14C). We speculated that the observed expression pattern of *NHX2* might be due to diurnal variation in expression of either miR158 or its target pseudo*-PPR*. However, though there was no variation in expression of miR158 at different time points of the day, the expression of pseudo-*PPR* was maximum at 8 hr followed by at 4 hr (Supplemental Figure 14D). Our results indicate diurnal expression pattern of both pseudo-*PPR* and *NHX2* might be due to diurnal variation of stoichiometric ratio between miR158 and pseudo-*PPR*.

### Down regulation of *NHX2* by PDP affects physiological processes

NHXs, mainly NHX1 and NHX2 are essential for turgor regulation and stomatal function (Andrés et al., 2014; Barragán et al., 2012). Therefore, to examine if PDP956-977 (+) enriched plants having altered *NHX2* expression also exhibit impaired stomatal functioning, we measured transpiration, stomatal conductance and water use efficiency of miR158 mutant, miR158 OE, pseudo-*PPR* mutant and pseudo-*PPR* OE lines at different time points of the day. All the lines showed reduced transpiration and stomatal conductance at different time points compared to the control, except miR158 mutant, whose transpiration rate was slightly higher at 8 hr light exposure and at dark (Figure 5C and 5D; Supplemental Dataset 9). Interestingly, pseudo-*PPR* OE also exhibited lower rate of transpiration and stomatal conductance as compared to the control, presumably due to higher stoichiometric ratio in favor of pseudo-*PPR* over miR158. Further, transpiration rate and stomatal conductance were maximum at 4 hr light and gradually decreased after 8 hr of light exposure in all but miR158 mutant and pseudo-*PPR* OE and finally all reached the lowest level at dark (after 2 hr 30 min dark). The impaired stomatal activity in miR158 mutant and miR158 OE was further examined by imaging the stomatal aperture at light and dark under scanning electron-microscope. It is evident that stomatal closure was delayed in miR158 mutant under dark as compared to control (Figure 5E; Supplemental Figure 15A). Also, miR158 mutant exhibited significantly higher water use efficiency (Figure 5F) and lower rate of water loss after 2 hr 45 min as compared to the control plants (Supplemental Figure 15B; Supplemental Dataset 9). Accordingly, when subjected to water-holding for 35 days, the miR158 mutant and pseudo-*PPR* OE could complete their life cycle, while the growth of the control Col-0, miR158 OE and pseudo-*PPR* mutant were severely affected and could not complete their life cycle (Figure 5G). Similar observations were recorded with NTL, ITL and DTL plants (Supplemental Figure 15C-15H). These results indicate biogenesis of PDP driven by miR158 regulate transpiration and stomatal conductance via *NHX2*.

Finally, we evaluated the effect of these altered physiological processes on plants growth and development by recording various life cycle and morphological parameters (Figure 5H; Supplemental Figure 16, Supplemental Dataset 9). Data suggested the miR158 mutant possessed elongated leaves and petioles with larger rosette area (Supplemental Figure 17) and bolted six days (± 1 day) earlier as compared to the Col-0 (Figure 5H). Bolting was delayed by 21days (± 2 days) in over expression lines than Col-0. Similarly, pseudo-*PPR* mutant matured 11 days (± 1 days) earlier than it’s over expression lines (Figure 5H and Supplemental Figure 16). Overall, both physiological and morphological data indicate miR158 regulate growth and development of *A. thaliana* by regulating *NHX2* via PDPs.

## DISCUSSION

### Natural variations in *MIR158* affect pri-miR158 processing

We identified natural variants of *MIR158* in west Himalayan populations of *A. thaliana*. The insertion/deletion events are unique to these populations, not observed in any other world accessions. Though both insertion and deletions were observed in these populations, notable is the discontinuous deletion one with very high frequency of occurrence (0.94) in a particular population (Mun). Subsequently, we identified some other populations besides Deh, Mun and Chit in the region and PCR screened the *MIR158* in these new populations as well. Interestingly, we could not identify any NTL plants in any of the populations including newly screened ones except, Chit and Deh. Noteworthy to mention here that all these populations are from same geographical continuity along the Himalayan elevation (700 m amsl to 2000 m amsl), except Chit which is at the extreme elevation of ∼3400 m amsl and lies hundreds of kilometer apart. It remains to be seen if abolition of miR158 activity in Mun population is under purifying selection aiding in local adaptation. Intuitively so, as there is gradual decrease in frequency of NTL plants as one moves from lower elevation (Deh), harboring almost equal frequency of NTL and ITL to higher elevation (Mun)that harbors high frequency of DTL plants (0.94) with no NTL plants.

The insertion/deletion altered the secondary structures of pri-miR158 (Figure 2D-2F) affecting their processing efficiency and/or accuracy by plant miRNA processing machineries, such as DCL1, HYL1, or SERRATE (Kurihara and Watanabe, 2004; Reinhart et al., 2002). Particularly, the positioning of the initial DCL1 processing event in many precursors is dependent on a lower stem of ∼15 bp below the miRNA/miRNA* duplex followed by a large internal loop (Cuperus et al., 2010; Mateos et al., 2010; Werner et al., 2010). These critical features of secondary structures, particularly the lower stem structure and internal loop is severely affected in DTL while in ITL, the internal loop is absent (Figure 2E and 2F). Thus the miRNA processing machineries completely or partially failed to recognize the inappropriate secondary structures (Figure 2E and 2F) resulting negligible or moderate expression in DTL and ITL, respectively(Figure 2A). Earlier, 139 bp insertion in primary transcript of miR163 in *A. arenosa* was shown to affect its processing leading to reduced expression (Ng et al., 2011). Different miRNAs may be processed by different mechanisms depending on the sequential direction of the miRNA processing machineries. miR158 is reported to be processed following canonical base-to-loop mechanism, where the first cut is made by counting a 15-nt stem region above the internal loop (Bologna et al., 2013). Overall, the insertion/deletion make secondary structures of pri-miR158 in appropriate for plant RNA processing machineries leading to reduced or negligible expression of miR158.

### Abolished miR158 activity triggers PDP biogenesis following “two-hit, two-cleavage” model

Earlier it was reported that miR158 targets a *PPR* and thereby regulates pollen development in *Brassica campestris* (Ma et al., 2017). However, our repeated attempts failed to identify the *PPR* as the target of miR158 in *A. thaliana*. Subsequently, based on our *in-silico* target prediction results, we validate a pseudo-*PPR* as its target. Though many pseudogenes are reported to generate siRNAs both in plants and animals (Guo et al., 2009; Thibaud-Nissen et al., 2009) and also act as sponge for miRNA (An et al., 2017; Yu et al., 2014), our findings that miR158 targets and cleaves a pseudogene may widen the range of biological functions of plant pseudogenes. Indeed, in absence of miR158 activity, the *TAS2* derived siRNAs, siR9 and siR12 bind to miR158 target pseudo-*PPR* and subsequently processed to phasiRNAs (Table 1). Thus, a large number of PDPs were generated in DTL and *MIR158* mutant plants as compared to NTL and ITL. Conversely, we detected negligible PDPs in pseudo-*PPR* mutant and miR158 OE lines. Although there is report that *TAS2*-derived siRNAs act as a trigger of tasiRNAs (e.g., tasiRNA2140) biogenesis from *PPR*, there is scanty of report showing cleavage of target by both the tasiRNAs for biogenesis of tertiary phasiRNAs (Chen et al., 2007; Howell et al., 2007). Moreover, considering the one-hit model is the most prevalent pathway for phasiRNA production and the two-hit model primarily reported from *TAS3*, the observed pathway of phasiRNA production implicates diversity in their biogenesis. More importantly, we observed phasiRNA production from both 5’ and 3’ cleaved ends of the target, albeit in higher number from 5’ cleaved end than 3’ cleaved end. Recently, similar two end target cleavage and phasiRNAs production from downstream of both cleavage sites were reported in wild strawberry (*Fragaria vesca*) *TAS3*S (Xia et al., 2015). The above study together with present report suggest in addition to prevalent “two-hit, one-cleavage” configuration for tasiRNA production (Liu et al., 2020), a “two-hit, two-cleavage’ configuration may be operating as well, at least in some plant species.

### A new module of phasiRNA regulation

Our observation that pseudogene act as a template for phasiRNA biogenesis implicates larger role of pseudogenes in plants. Firstly, pseudogenes are known template for 24-nucleotide siRNA biogenesis, primarily involve in RNA directed DNA methylation (Guo et al., 2009). In contrast, we observed a large number of 21-nucleotide PDPs those were associated with AGO1 (Supplemental Figure 11b and Supplemental Data 6) indicating probable involvement in cleavage mediated function. Secondly, pseudogene derived siRNA targeting of protein coding gene is not known in plants. Here, we show that one of the PDPs targets *NHX2*, a member of *NHX* gene family involve in vacuolar K^+^ homeostasis (Barragán et al., 2012). Further, in other organisms most of the pseudogene derived siRNAs are known to target their parent protein coding genes (Tam et al., 2008; Watanabe et al., 2008) rather than non-parental gene. The PDPs neither showed significant similarity with parent *PPR* (AT1G64100) nor any *PPR* target could be identified in degradome data. The targeting of *NHX2* by one of the PDPs also contradicts earlier report that opined against secondary targeting by clad-derived siRNAs on non-clad target transcripts, especially in case of *PPR* derived phasiRNAs (Howell et al., 2007). Noteworthy to mention that phasiRNAs production is restricted only to P-type *PPR* in plants (Howell et al., 2007; Xia et al., 2013) and phylogenetic analysis of the template pseudo-*PPR* shows it belongs to P-type PPR clad (Supplemental Figure10). Secondary siRNA biogenesis pathway involving PPRs following the miRNA-*TAS* like gene-*PPR*-siRNA module (Xia et al., 2013) or the most common pathway i.e miRNA-*TAS*-siRNA are mainly triggered by 22-nucleotide siRNAs (a few are of 21-nucleotide e.g., miR156 and miR172) following “two hit, one cleavage” model (Zhai et al., 2011). We demonstrate another layer of 21-nucleotide phasiRNA biogenesis, miRNA-*TAS*-siRNA-pseudo-*PPR*-tertiary phasiRNA-*NHX2* that regulates a protein coding gene of far distant family following “two hit, two cleavage” model (Figure 6).

**Fig. 6:**
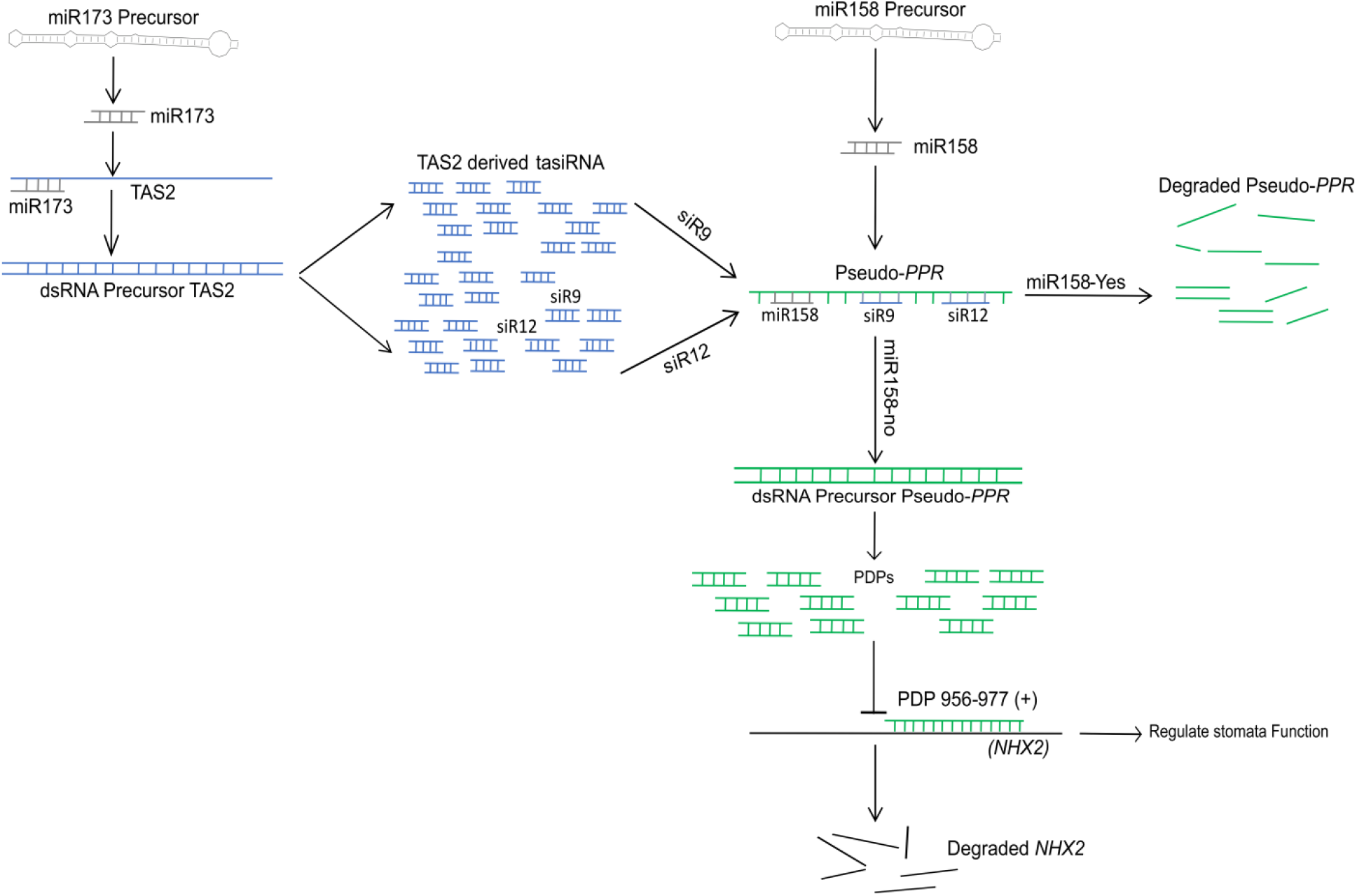
Proposed model for miR158 regulation. The model depicts miR158 acts as a negative regulator of phasiRNA biogenesis from a pseudo-*PPR*. The pseudo-*PPR* is targeted by miR158 besides being targeted by two tasiRNAs, siR9 and siR12 derived from *TAS2* locus. While miR158 activity leads to degradation of the pseudo-*PPR* and thus there is no biogenesis of phasiRNAs, null activity of miR158 makes the pseudo-*PPR* template available for targeting by the two tasiRAs leading to biogenesis of phasiRNAs. One of the phasiRNA produced by this mechanism targets and degrades the *NHX2* gene which in turn regulates stomatal functioning.

### PDP mediated down-regulation of *NHX2* affects physiological processes in *A. thaliana*

The down-regulation of *NHX2* by a PDP (Figure 5B) led to reduced transpiration, stomatal conductance and better water use efficiency (Figure 5C-5G). The reduced transpiration and stomatal conductance in miR158 mutant and pseudo-*PPR* OE as compared to control is in accordance with earlier observation of NHX2 mediated regulation of these activities (Andrés et al., 2014). The delayed response of stomatal functioning has been attributed to NHX proteins which help to maintain turgor by regulating K^+^ homeostasis and it corroborates with the observed diurnal variations in *NHX2* expression in different lines (Supplemental Figure 14D). Both opening and closure of stomata was reported to be affected in double knock out mutant of *NHX1* and *NHX2* in *A. thaliana* and the single mutant was more affected mainly in stomatal opening than closure (Andrés et al., 2014). It is evident that the stomatal opening was more impaired than closure in miR158 mutant as compared to other lines, as the rates of transpiration and stomatal conductance did not fall rapidly as others as dusk approached. In fact response to stomatal closure was delayed showing higher rates at dusk (8 hr) and even at dark (2hr 30 min). The reduced transpiration could not be attributed to the defective guard cells (Figure 5E). Notably, miR158 mutant and to a lesser extent pseudo-*PPR* OE lines completed life cycle earlier than control both under water stress as well as under regular water supply conditions (Figure 5G and 5H). The overall performance of pseudo-*PPR*OE is in line with the observed phasiRNA generation and probable stoichiomeric ratio between pseudo-*PPR* and miR158 in favor of pseudo-*PPR* in this line. Thus the negligible expression of miR158 and enhanced phasiRNA generation might be such a strategy of the population thriving under elevated area to complete life cycle early.

Natural variants of *MIR158* led us to identify a module of 21-nucleotide phasiRNA biogenesis. The identified pathway widens the scope of phasiRNA biogenesis beyond prevalent pathways. miRNA driven cleavage of a pseudogene or regulation of a protein coding gene by a pseudogene derived phasiRNA highlights the significance of pseudogene as well as role of tertiary phasiRNA in plants. However, although siR9 and siR12 act as a trigger for biogenesis of phasiRNA, it will be interesting to examine if a 21- or 22-nucleotide miR158 can act as a trigger for observed phasiRNA biogenesis from the cleaved end instead of template degradation.. Understanding the evolutionary significance of such a pathway may shed light on siRNA regulated local adaptation.

## METHODS

### Plants materials and growth conditions

The seeds of different populations of *A. thaliana* of west Himalayas (Deh, Mun, Chit), and SALK mutants lines (miR158 mutant, SALK_025691; pseudo-*PPR* muatnt, SALK_043644) and Col-0 were grown under controlled condition of 22°C temperature, 140 µmol/m^2^/sec light intensity and 16/8-hour day/night cycle. Besides, seeds were also grown in common garden at CSIR-NBRI, Lucknow, Uttar Pradesh, India during the month of November to February to check the influence of environment on the expression of miR158. The homozygous lines of SALK mutants were selected as discribed in GABI (http://signal.salk.edu/tdnaprimers.2.html) using primers listed in Supplemental Dataset 10.

### RNA isolation, small RNA sequencing and analysis

Total RNA was isolated from rosette leaves of 28 days old plants using mirVana™ miRNA isolation kit (Ambion, USA) following the manufacturer instructions. The quality of RNA was checked on both agarose gel as well as using bioanalyzer (Agilent, USA). RNA samples having RIN >7 were used for small RNA library preparation. Four pooled small RNA libraries corresponding to Deh, Mun, Chit and Col-0 was prepared and used as replicates of earlier sequenced data from the same populations (Tripathi et al., 2019). Further, for each plant type individual libraries were prepared and sequenced in replicates, except for DTL, miR158 OE and pseudo-*PPR* OE (one of these libraries failed the quality control).Libraries were prepared using NEB Next Small RNA Library Prep Set (New England Biolabs) and sequencing was carried out using Illumina HiSeq 4000 in single-end mode by outsourcing. Thus, a total of 20 small RNA sequencing libraries data were analyzed (Supplemental Dataset 11).The raw sequences were analyzed using UEA small RNA workbench (Version 4.5) (Stocks et al., 2018). Briefly, the adapters were removed from raw reads and size-filtered between 18-28 nucleotides using adapter removal tool. The filtered reads were mapped onto the reference *Arabidopsis* TAIR10 genome using Bowtie2 with default parameters. The genome mapped reads were used for identification and quantification of miRNA and tasiRNA using miRProf implemented in UEA workbench with miRBase22 (http://www.mirbase.org/) and tasiRNA data base (http://bioinfo.jit.edu.cn/tasiRNADatabase) (Zhang et al., 2014), as background, respectively.

### PhasiRNA identification and scoring

For identification of pseudo-PPR derived phasiRNA (PDP), the filtered small RNA reads were mapped onto the pseudo-PPR reference sequence using Bowtie2.0 (zero mismatch) (Langmead and Salzberg, 2012). It is worthy to mention that binding site of one of the *TAS2* derived small RNA, siR9 on the pseudo-*PPR* and its production site on *TAS2* show perfect complementary to each other (Supplemental Figure 17A). Therefore, in order to exclude any ambiguous mapping of siR9 (higher expression as compared to PDPs) on pseudo-*PPR*, we masked (replaced by N) half complementary sites of pseudo-*PPR* (Supplemental Figure 17B). This led us to specifically identify the PDPs.

To identify whether this pseudo-*PPR* acts as true phasi generating loci, the p value was calculated using the tasi-predictor tool in-built in UEA workbench and pseudo-*PPR* sequence as reference following (Chen et al., 2007). Further, the phasing score was also calculated using 21-nucleotide phasing register in a 10 cycle window and p > 0.001. Phasing score was calculated following (De Paoli et al., 2009).

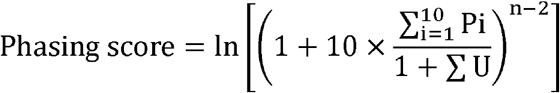

where n is the number of phase registers occupied by at least one unique 21-nucleotide sRNA within a ten-phase register window, P is the total number of reads for all 21-nucleotide sRNA reads falling into a given phase in a given window, and U is the total number of unique reads for all 21-nucleotide sRNAs falling out of a given phase.

### Promoter analysis and identification of polymorphism in *MIR158*

The putative promoter sequences of *MIR158* (one kb upstream of transcription start sites) were extracted from our genome sequence data of Deh, Mun, and Chit and were checked for any sequence variations using multiple sequence alignment tool implemented in MEGA 6.0 (Tamura et al., 2013). During initial PCR amplification of *MIR158* locus from a few plants of the three populations, we detected length polymorphisms as visualized on 1.2% agarose gel which was confirmed by sequencing of the amplified products. Subsequently, *MIR158* locus was PCR amplified and sequenced from 50 plants of each of the three populations using primers listed in Supplementary Data 10. Multiple sequence alignment analysis was performed using MEGA 6.0. To further confirm the length polymorphisms, we amplified the pri-miR158 locus using cDNA as template from different populations using the length polymorphisms specific primers (NTL, ITL and DTL specific primers, Supplementary Data 10). The miR158 locus of these populations were compared with 1135 *A. thaliana* accessions as well using POLYMORPH 1001 tools (https://tools.1001genomes.org/polymorph/) selecting nucleotides position range 3365982-33666000 on chromosome 3 (range of miR158a locus).

### Secondary structure prediction of pri-miR158

The primary transcript of miR158 amplified from each of the three populations were sequenced and secondary structure of these primary transcripts were predicted using RNAfold web server (http://rna.tbi.univie.ac.at/cgi-bin/RNAWebSuite/RNAfold.cgi) at default parameters.

### Target prediction and degradome analysis

Targets of miRNA, tasiRNA and phasiRNA were predicted by psRNA target (Dai and Zhao, 2011) using expectation value cutoff of less than three. Available degradome data of *A. thaliana* (GSE55151) (Thatcher et al., 2015) was downloaded and analyzed using PAREsnip (Folkes et al., 2012) following stringent criteria. Briefly, mismatch on position 10 and 11 were not allowed, maximal numbers of mismatches were set to four and two mismatches on adjacent positions were not allowed. The targets with lower confidence (category four and three) were not considered. The T-plots were generated by VisSR tool (Stocks et al., 2018).

### Target validation by modified 5’ RLM-RACE

To validate the putative targets of miRNA and phasiRNA, modified 5’RLM -RACE was carried out using GeneRacer^™^ Kit (Ambion, USA) following user’s manual. Briefly, RNA was isolated using RNeasy Mini Kit (Qiagen, USA). Adapter was ligated to 4µg freshly isolated RNA at 37°C for one hours followed by 16 °C for eight hours. cDNA was synthesized using random primers with ligated RNA. Touchdown PCR amplification was performed using the adapter specific GeneRacer 5’ primer and gene specific outer primers following PCR program of 2 min at 94°C, followed by five cycles of 30s at 94°C and 90s at 72°C, five cycles of 30s at 94°C and 120s at 70°C, and 25 cycles of 30s at 94°C, 30s at 65°C and 90s at 72°C and finally 10 min at 72°C. Nested PCR amplification was performed using the adapter specific GeneRacer 5’ Nested primer and gene specific inner primers and two microliter template from the initial touchdown PCR following the PCR program of 2 min at 94°C, followed by 25 cycles of 30s at 94°C, 30s at 65°C and 2 min at 72°C and finally 10 min at 72°C. The PCR product was run on 2% agarose gel and the expected bands were eluted using Wizard^®^ SV Gel and PCR cleanup kit (Promega, USA),cloned in pGEM-T easy vector (Promega, USA), and at least five individual clones were sequenced for each target.

### Expression analysis by qRT-PCR

Expression level of miRNA158 primary transcripts, precursor and target genes were carried out using qRT-PCR. cDNAs were prepared by GoScript™ Reverse Transcription System following manufactuere’s protocol (Promega, USA). All qRT-PCRs were performed in ABI 7300 real-time PCR machine (Applied Biosystem, USA) using DyNAmo Flash SYBR Green (Thermo, USA) with cycling conditions: denaturation at 95°C for 10 min followed by 40 cycles of denaturation at 95°C for 20s, annealing and extension together at 60°C for 60s. The respective primers are listed in Supplemental Dataset 10. All reactions were performed with three to five biological and two technical replicates. 5s-rRNA and actin gene were used as endogenous control for stem loop-PCR and target genes, respectively (Czechowski et al., 2005; May et al., 2013). The expression value of each sample was normalized using expression value of respective endogenous control and relative expression was calculated. The fold change values were determined using 2^−ΔΔCt^ method (Schmittgen and Livak, 2008). Before conducting stem loop qRT-PCR, the specificity of miR158 stem loop primers was checked using the end point PCR following (Varkonyi-Gasic et al., 2007). Briefly, 100 ng of total RNA was revesre transcribed using miR158 stem loop primers and SuperScript® III Reverse Transcriptase (Invitrogen, USA) following manufacturer’s instructions. The polymerase chain reaction was performed using two µl of cDNA as template and miR158 specific forward primer, and stem-loop specific reverse primer at thermocycler condition of 94°C for 2 min, followed by 35 cycles of 94°C for 15s and 60°C for 1 min. The PCR product was checked on 2% agarose gel (Supplemental Figure 18).

The variation in expression of the primary transcripts of miR158a and miR158b was taken as proxy to distinguish between the expression of miR158a and miR158b. This is because the precursor sequences of miR158a and miR158b do not harbour sufficient variations (Supplementary Fig.1b).

### Phylogenetic analysis of PPRs

297 PPRs having conserved PPR-domains amongst 450 reported *A. thaliana* PPRs(Lurin et al., 2004) were considered for phylogenetic analysis. The sequences of the PPRs were downloaded from TAIR (www.arabidopsis.org) and aligned using CLUSTAL-W implemented in MEGA 6.0 (Tamura et al., 2013). Phylogenetic tree was constructed using Neighbor-Joining method implemented in MEGA 6.0 with 1000 bootstrap replicates.

### Cloning and generation of over expression transgenic plants

Pri-miR158 and pseudo-PPR fragments were PCR amplified using respective gene specific primers (Supplemental Dataset 10) following standard PCR protocol. The amplified fragments were cloned in pGEM-T cloning vector (Promega, USA) and subsequently transferred into pBI121 binary vector. *A. thaliana* plants (Col-0 background) were transformed by floral dip method (Clough and Bent, 1998). Transformants were screened on one-half-strength MS agar plates containing kanamycin (50µg/mL) followed by PCR screening. Subsequently, T3 homozygous plants were selected for further analyses. In spite of repeated attempts we were unable to generate the complementation line of miR158 in DTL plants due to difficulty in getting viable seeds after transformation due to unknown reasons.

### Measurement of morphological and physiological parameters

Various morphological and life cycle parameters of ten five-weeks post-germinated plants each of miR158 mutant, pseudo-*PPR* mutant, transgenic lines and Col-0 were measured. Leaf area was estimated as product of length and width. Leaf shape was represented as a ratio of length to width. Rosette area was measured as product of major and minor axis and rosette shape as the ratio of major to minor axis. Both leaf and rosette shape were considered as rounded when the ratio is close to one and elongated if the ratio was more than one. Life cycle namely, bolting, first flower opening, emergence of first silique and senescence were measured in terms of days after germination. Plants were monitored regularly and data were recorded at every two alternate days.

Stomatal conductance, rate of transpiration, and water use efficiency of plants at bolting stage (principal growth phase 5.10 (Boyes et al., 2001))were measured using LICOR-6400 portable photosynthesis system (LI-COR, USA), at four different time points: Immediately after onset of light, four and eight hours post exposure of light and two and half hours post exposure of dark. Stomatal aperture was measured by peeling the epidermal layer of leaf lower surface and mounted in glycerol and visualized under light microscope at 100X using LEICA DM2500. Stomatal measurement was taken at two time points of four hours post light and two and half hours of post dark exposure. Additionally, the stomata from these two time points were also visualized and measured using scanning electron microscopy (FEI Quanta 250, Bionand, Spain). The leaf tissues were fixed and processed according to (Bomblies et al., 2008). Briefly, the tissue was fixed overnight at 4°C in FAA fixative (3.7% formaldehyde, 50% ethanol, 5% acetic acid) and then dehydrated serially in the ethanol solutions of 30%, 50%, 70%, 90%, 95% and 100% for about 1hour each. The tissue was kept at 4°C overnight in 100% ethanol and then processed for CPD (Critical-Point Drying) followed by gold coating and visualization at 8000x.

### Drought Assay

Ten plants of each of miR158 mutant, miR158 OE, pseudo-*PPR* mutant, pseudo-*PPR* OE and Col0 along with NTL, ITL and DTL plants were grown under controlled condition for three weeks with normal watering. On the last day of third week pots were saturated with water and soil surface was covered with plastic wrap. The plants were allowed to grow without additional water and monitored till they survived. The photographs were taken after 35 days of water holding.

### Relative Water Content

The relative water content and water loss was measured following (Barrs and Weatherley, 1962). Briefly, rosette leaves were harvested and soaked in water for 30 min to make them completely turgid. The turgid weight of each leaf was recorded and then kept at room temperature for varying intervals of time. The weight of each leaf was measured after 5min, 15min, 30 min, 1 hour, 1.45 hour and 2.45 hour. After 2.45 hour, the leaves were kept at 65°C in dry oven for 72 hours and then measured the dry weight of the leaves. The relative water content was measured using the formula:

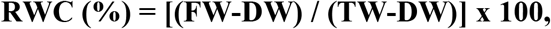

Where, RWC = relative water content, FW= fresh weight at different time points, DW= dry weight, and TW=turgid weight

All the morphological and physiological data was analyzed using one-way ANOVA in R. The multiple comparison analysis of the linear ANOVA model was conducted using ‘glht’ function in multcomp package in R.

## Supplemental Data

**Supplemental Figure 1**. Sequence conservation of miR158 (Supports **Figure 1)**

**Supplemental Figure 2**. Relative Expression of miR158 in different populations as estimated by stem-loop RT-PCR (Supports **Figure 1)**

**Supplemental Figure 3**. Multiple sequence alignment of miR158 promoter sequences from different populations using CLUSRTAL-W (Supports **Figure 1)**

**Supplemental Figure 4**. Representative image of PCR amplification of miR158 locus showing length polymorphisms in different populations of Indian west Himalayas (Supports **Figure 1**)

**Supplemental Figure 5**. Length polymorphism of miR158 loci in different populations (Supports **Figure 1**)

**Supplemental Figure 6**. Length polymorphism in pri-miR158 locus in NTL, DTL and ITL plants (Supports **Figure 1)**

**Supplemental Figure 7**. Predicted secondary structures of miR158 precursor (Supports **Figure 2)**

**Supplemental Figure 8**. Target plot (t-plot) for miR158 and tasiRNA targets identified using degradome data (Supports **Figure 3**)

**Supplemental Figure 9**. Translated protein sequence of pseudo-*PPR* with predicted ORFs (open reading frames) (Supports **Figure 3**)

**Supplemental Figure 10**. Phylogenetic analysis of *PPR* genes (Supports **Figure 3**)

**Supplemental Figure 11**. Abundance of phasiRNA (PDPs) in different populations (Supports **Figure 4**)

**Supplemental Figure 12**. Differential expression of PDPs using log2 values of read counts from different plant samples (Supports **Figure 4**)

**Supplemental Figure 13**. Target-plot (T-plot) of *NHX2* (Supports **Figure 5**)

**Supplemental Figure 14**. Relative expression pattern of *NHX2, NHX1*, pseudo-*PPR* and miR158 using qRT-PCR and stem-loop qRT-PCR (Supports **Figure 6**)

**Supplemental Figure 15**. Physiological parameters of different samples (Supports **Figure 6**)

**Supplemental Figure 16**. Growth and leaf size of different transgenics and Himalayan populations (Supports **Figure 6**)

**Supplemental Figure 17**. Morphological variations in different transgenic lines and Col0 (Supports **Figure 6**)

**Supplemental Figure 18**. Masked sequence of pseudo-*PPR* used for identifying pseudo-*PPR* derived phasiRNAs (PDPs) (Supports **Figure 4**)

**Supplemental Figure 19**. Gel Image of Endpoint PCR in the three populations (Supports **Figure 1**)

**Supplemental Table 1**. Length distribution of expressed form of siR9 and siR12 in NTL, ITL and DTL.

**Supplemental Dataset 1**. Expression (read counts) of miRNAs, siR9 and siR12in different populations of Indian west Himalayas

**Supplemental Dataset 2**. Expression (read counts) of miRNAs, siR9 and siR12 in NTL, ITL and DTL.

**Supplemental Dataset 3**. List of putative targets of miR158 as predicted using psRNATarget.

**Supplemental Dataset 4**. Pseudo-*PPR* derived phasiRNA (PDPs) and their expression.

**Supplemental Dataset 5**. Comparison of phasiRNA derived from 23 *A. thaliana PPRs* and the PDPs along with their read counts and sequence.

**Supplemental Dataset 6**. AGO1 and AGO4 associated PDPs.

**Supplemental Dataset 7**. Significant small RNA cluster (p value <0.01) on pseudo-*PPR*. Table also includes the phasiRNA and their expression in different plants identified from highly significant cluster (Start 746-End 997).

**Supplemental Dataset 8**. List of identified targets of PDP 956-977 (+) using PAREsnip tool.

**Supplemental Dataset 9**. Test of significance of variation (ANOVA) for the measured physiological and morphological traits along with pair-wise comparison.

**Supplemental Dataset 10**. List of primers and their sequences used in the study.

**Supplemental Dataset 11**. Sequencing and mapping read statistics of the small RNA libraries analyzed in the study.

## ACKNOWLEDGEMENTS

The authors acknowledge the financial support of the Department of Biotechnology (DBT), (grant # BT/PR23518/BPA/118/236/2017), Government of India, and partly by Council of Scientific and Industrial Research (CSIR), New Delhi, India. AKV also acknowledge the University Grant Commission (UGC), New Delhi for providing the fellowship. The authors are thankful to in-charge, scanning electron microscope, central instrumentation facility at CSIR_NBRI, Lucknow, UP, India. The institutional ethic committee reference number for the manuscript is CSIR-NBRI_MS/2020/11/03.

## AUTHOR CONTRIBUTIONS

AMT conducted experiments, analyzed the data and drafted the MS; RS, AKV and PM conducted experiments; AS analyzed the data and drafted the MS; SN and PAS helped and conducted physiological experiments; SR designed experiments, analyzed data and wrote the MS.

