## Supplementary Information for "Abolished miR158 activity leads to 21-nucleotide tertiary phasiRNA biogenesis that targets *NHX2* in *Arabidopsis thaliana*"

Supplemental Figures 1 to 19

Supplemental Table 1

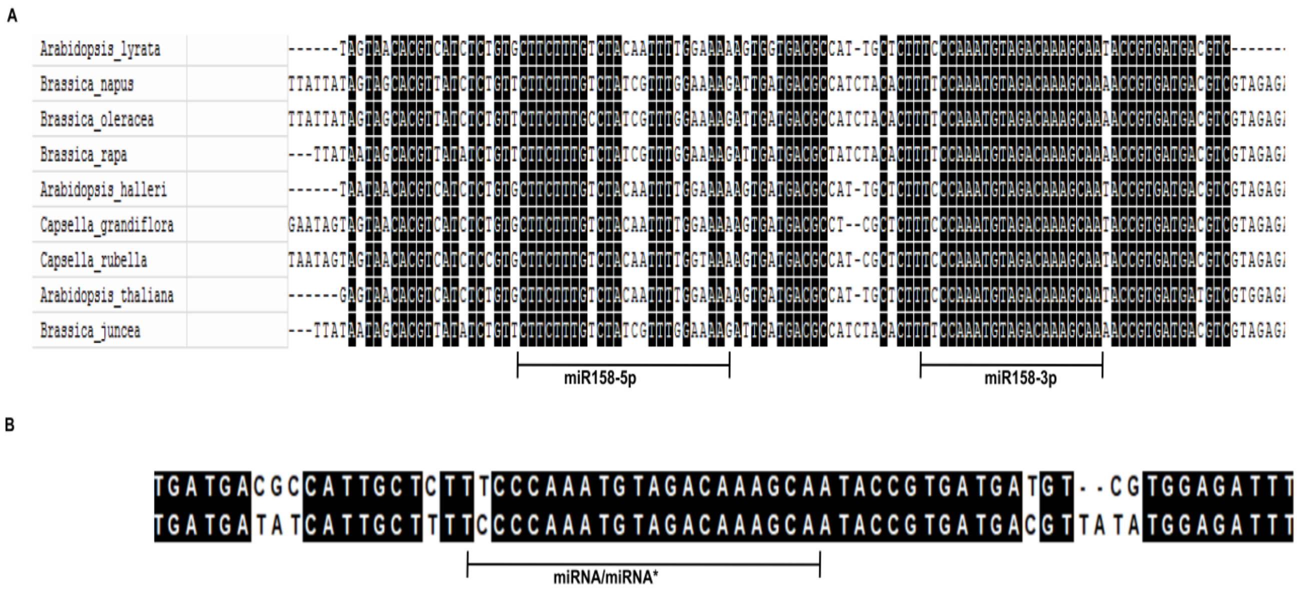


**Supplemental Figure 1. Sequence conservation of miR158**

**(A)** Multiple sequence alignment of miR158 loci from different members of the brassicaceae family. The dark shaded regions show the conserved sites among the members. The horizontal bar indicates the sequence of 5p and 3p arms of miR158.

**(B)** Similarity between precursor sequence of miR158a and miR158b in *Arabidopsis thaliana*. The dark shaded region indicates the conserved sites with horizontal bar indicating miRNA and miRNA star sequences.

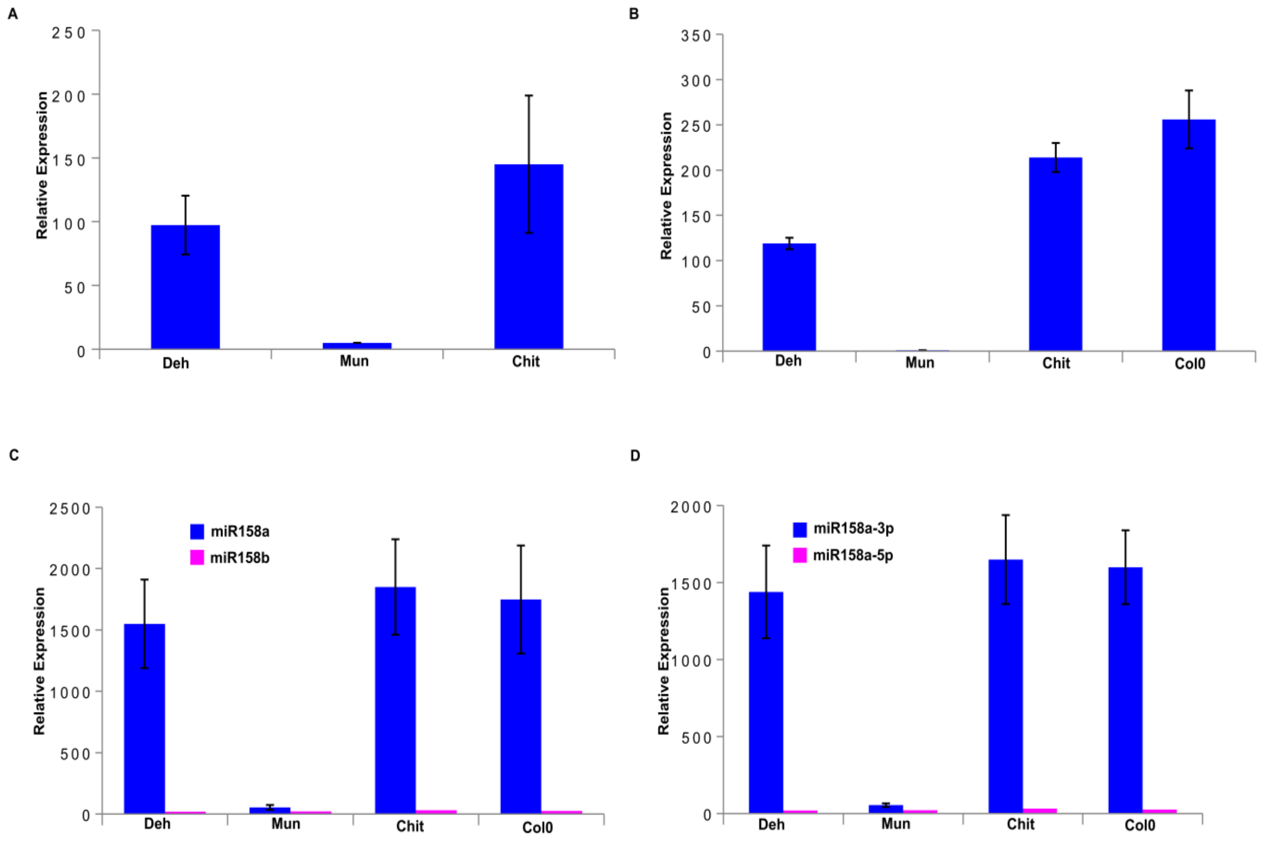


**Supplemental Figure 2. Relative Expression of miR158 in different populations as estimated by stem-loop RT-PCR**

Expression pattern of miR158 under: **(A)** native field (west Himalayas); and **(B)** common garden conditions. The data represents the mean of five independent biological replicates and line above bars represents ± SD.

Comparative expression pattern of: **(C)** miR158a and miR158b; and **(D)** 158a-3p and 158a-5p. The data represents the mean of three independent biological replicates and line above bars represents ± SD.

The expression data was normalized by the expression of 5s0rRNA endogenous control and relative expression was calculated using $2^{-\Delta\Delta Ct}$ method

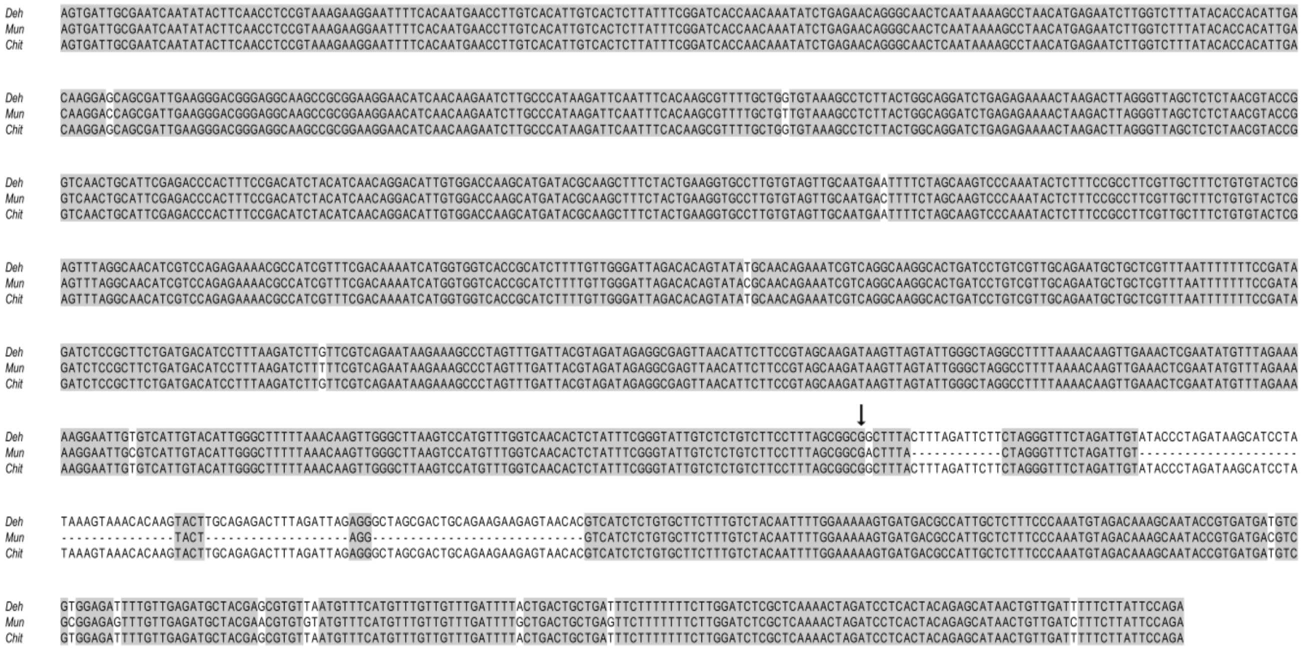


**Supplemental Figure 3. Multiple sequence alignment of miR158 promoter sequences from different populations using CLUSTAL-W**

One kb upstream sequence of TSS of miR158 was extracted from the three previously sequenced genomes of the three populations (unpublished) as putative promoter regions. The grey shaded region indicates the sequence conservation and the Transcription Start Site (TSS) of miR158 gene is marked by an arrow. Sequences downstream of the TSS are the miR158 gene sequences of the three populations showing the polymorphism at the miR158 loci.

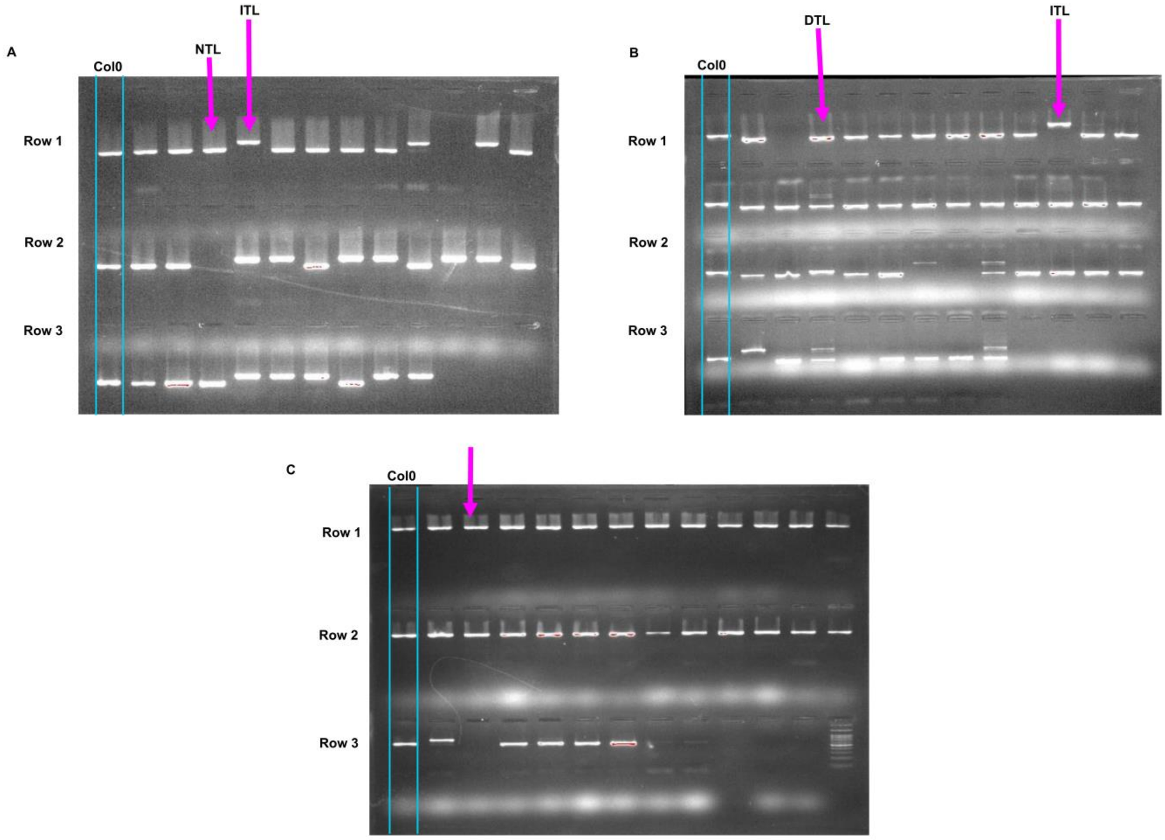


**Supplemental Figure 4. Representative image of PCR amplification of miR158 locus showing length polymorphisms in different populations of Indian west Himalayas**

Variations in the size of miR158 locus in different individuals of **(A)** Deh; **(B)** Mun; and **(C)** Chit populations. The first lane (marked with cyan) represents the length of miR158 locus in Col0 (Normal type locus, NTL); ITL, Inserted type locus and DTL, Deleted type locus (indicated by arrow).

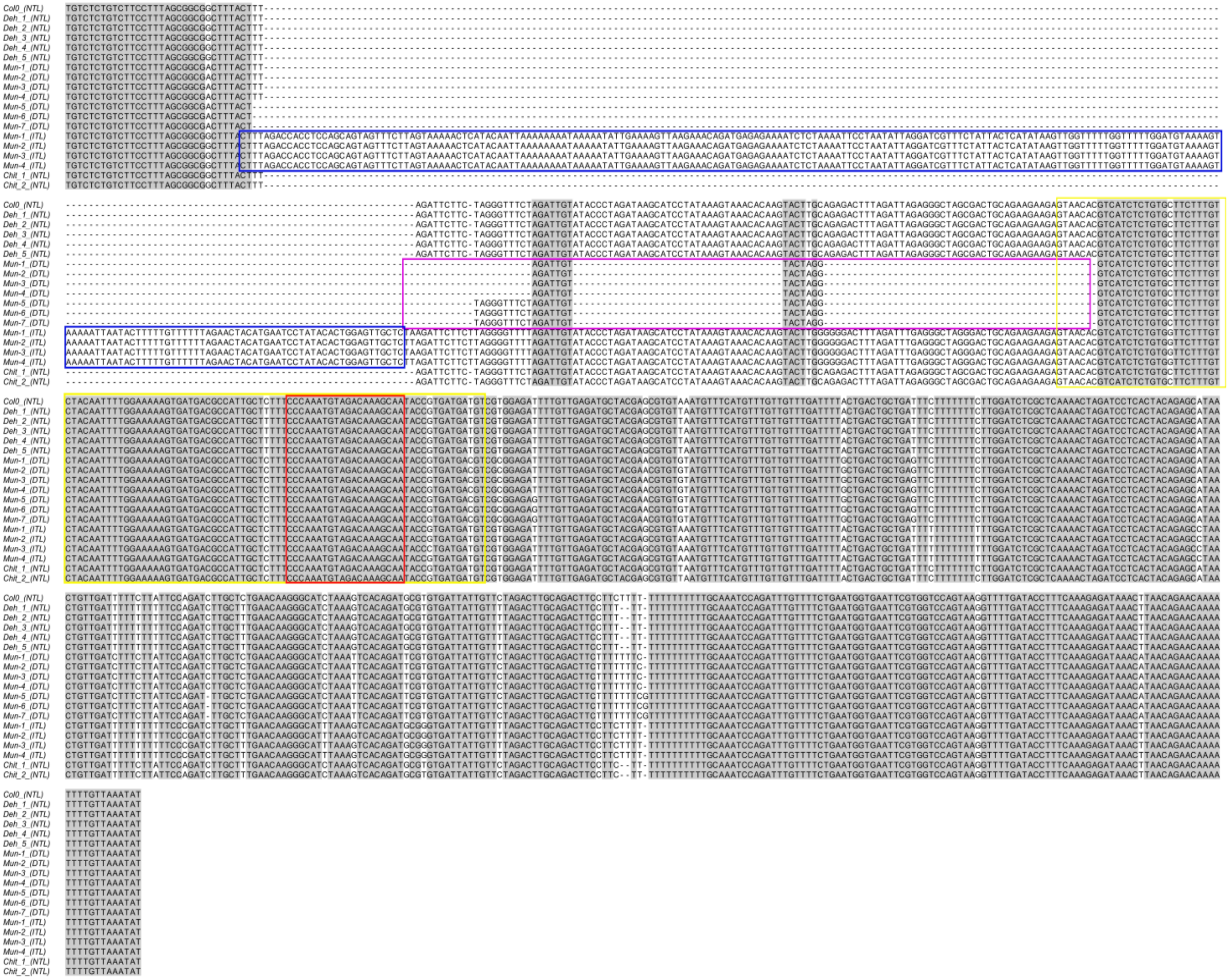

**Supplemental Figure 5. Length polymorphism of miR158 loci in different populations**

Multiple sequence alignment of miR158 loci amplified and sequenced from different west Himalayan populations along with Col0 is shown. The insertion sequence in ITL and the deleted sequence in DTL are shown in blue and violet box, respectively. The precursor and mature sequences of miR158 are shown in yellow and red boxes, respectively. The conserved regions are shaded in grey.

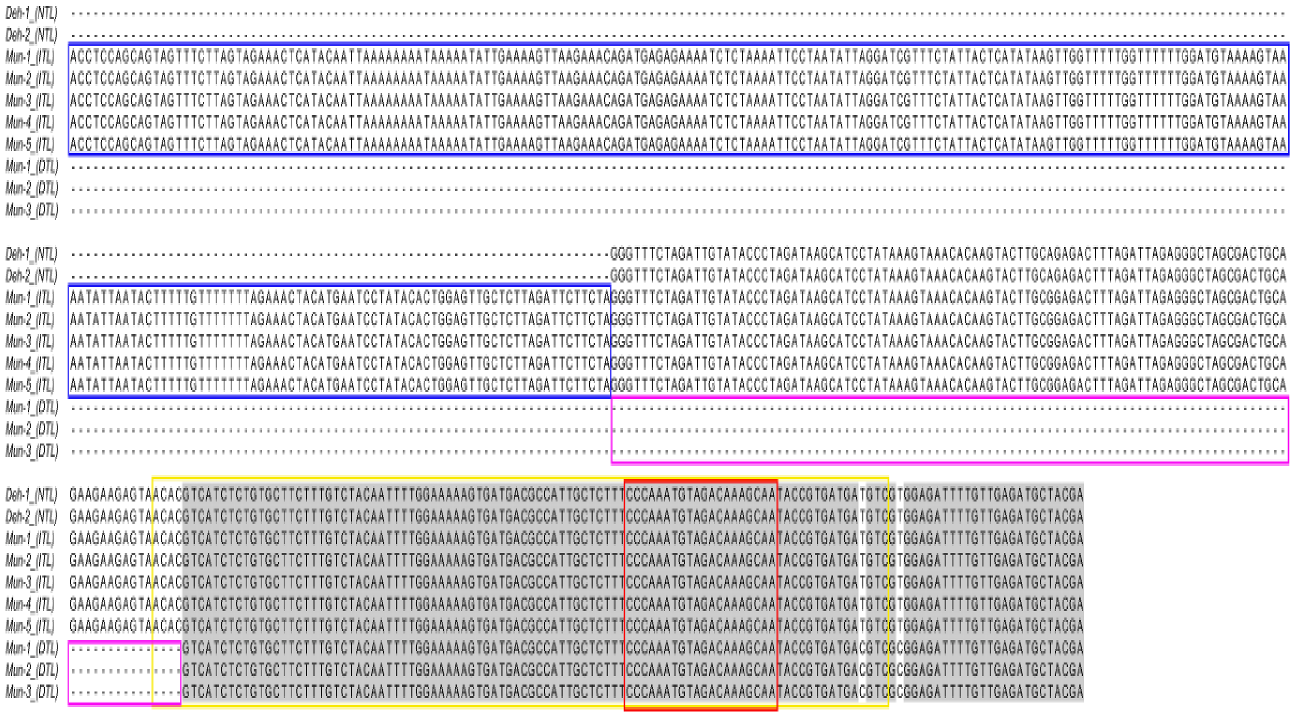

**Supplemental Figure 6. Length polymorphism in pri-miR158 locus in NTL, DTL and ITL plants**

Multiple sequence alignment of pri-miR158 sequenced from cDNA as template from ITL, DTL and NTL plants. The insertion sequence in ITL and the deleted sequence in DTL are shown in blue and violet box, respectively. The precursor and mature sequences of miR158 are shown in yellow and red boxes, respectively. The conserved regions are shaded in grey.

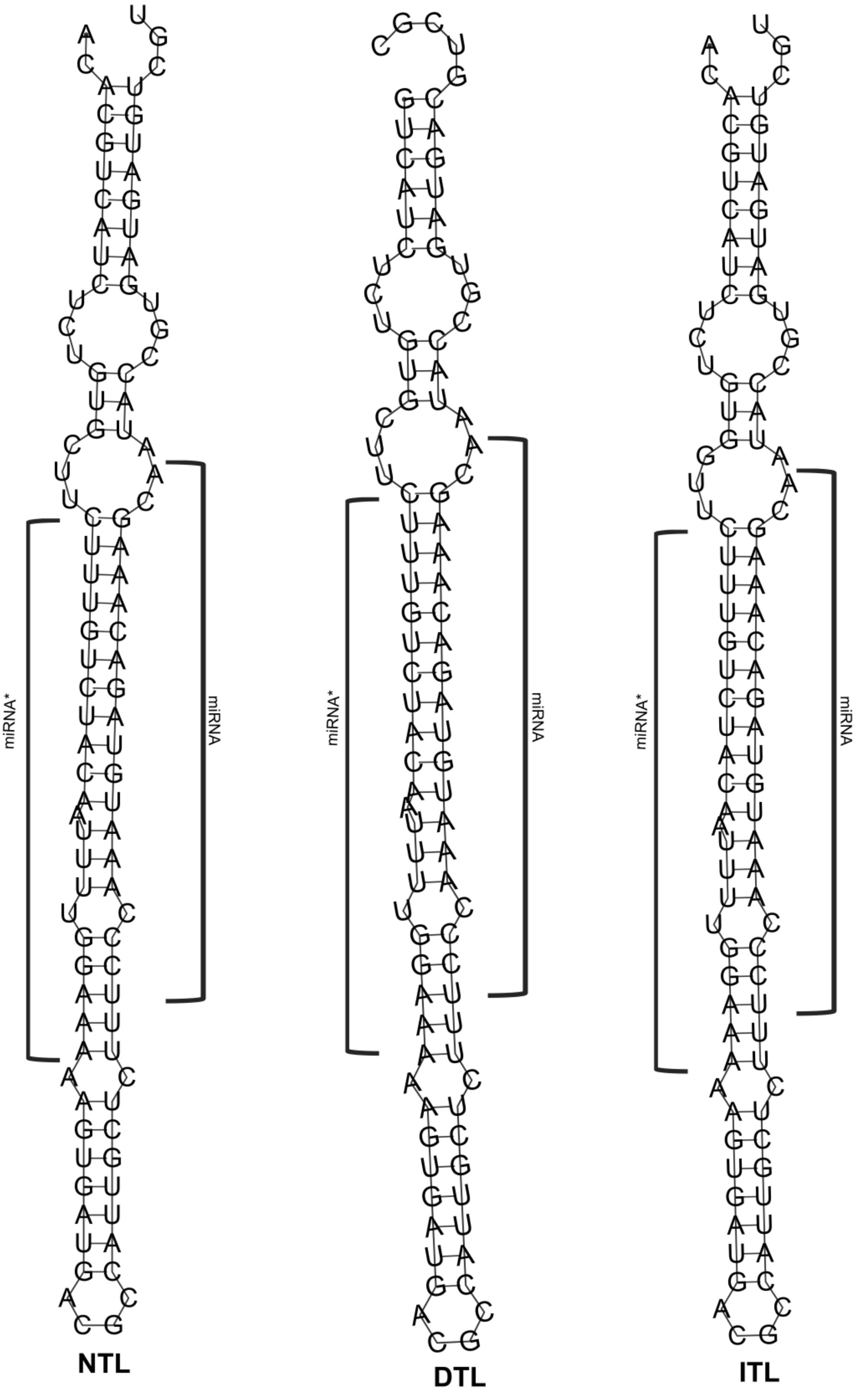


**Supplemental Figure 7. Predicted secondary structures of miR158 precursor**

The secondary structure of miR158 precursors in (a) NTL, (2) DTL and (c) ITL. The miRNA and miRNA star (*) sequence are shown in square brackets. The precursor sequence was annotated using pre-miR158 sequence reported in miRBase20.0 and the secondary structures were predicted using RNAfold.

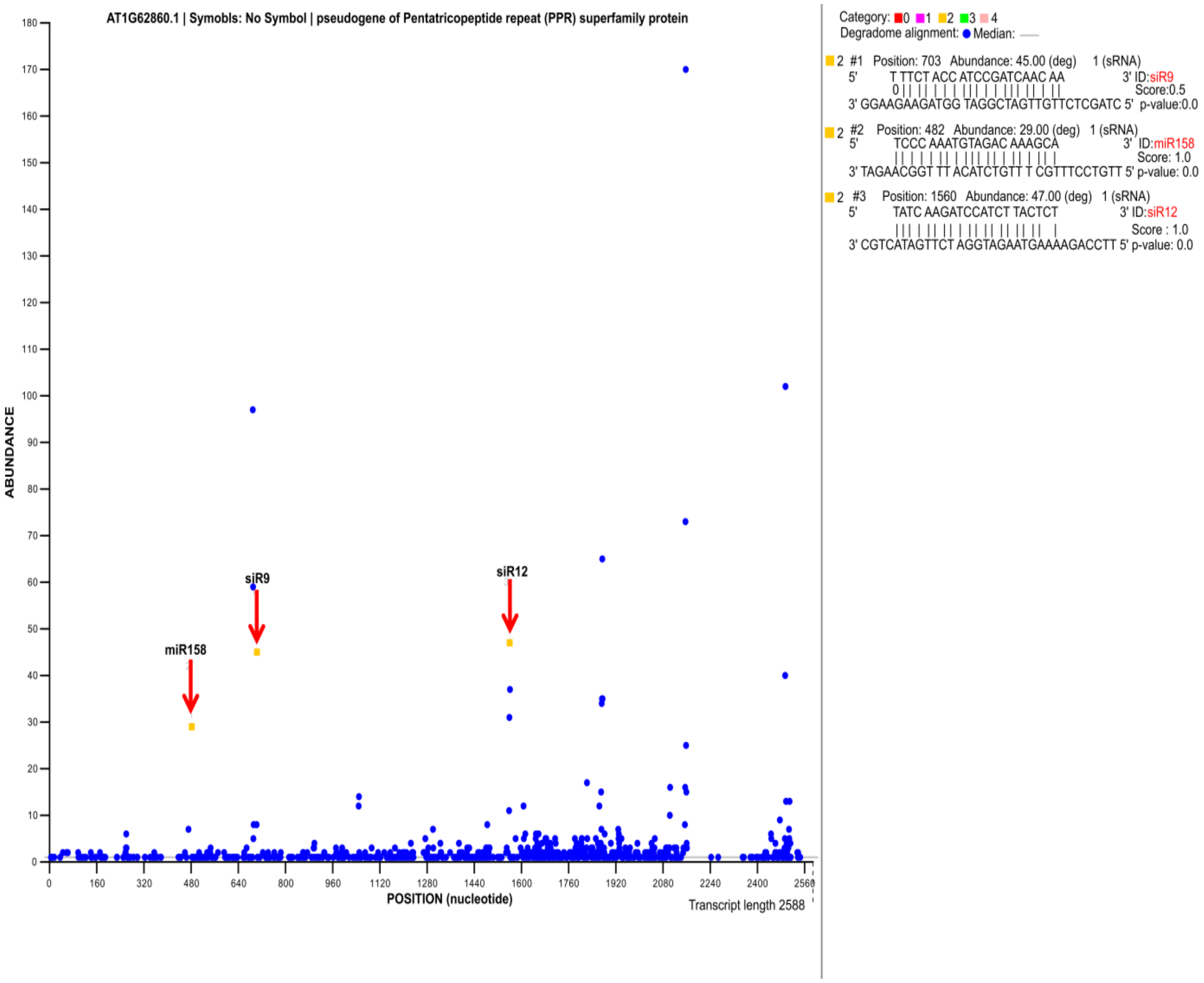


**Supplemental Figure 8. Target plot (t-plot) for miR158 and tasiRNA targets identified using degradome data (GSE55151)**

Target plots (t-plots) of miR158 and tasiRNAs- siR9 and siR12 using degradome data. Red arrows (Orange Square) indicate signatures consistent with miRNA and tasiRNA directed cleaved sites of pseudo-*PPR*. The blue circles represent the fragments mapped on the pseudo-*PPR*. The X and Y axis represents nucleotide position of pseudo-*PPR* and normalized read abundance (TPM, transcripts per million) of cleaved transcript detected in degradome-seq, respectively. mRNA:miRNA/tasiRNA alignments along with p-value, alignment score and cleavage positions are also shown.

5'-3'Frame

APPFVVVRV**Stop**RQRKSGESKMetLARVCRSESSSGNAAAVSARLFSTRLVHRRVA
KKTSKVEKSMetPSLRSF**Stop**RREFEIEKWISLYQKLR**Stop**CDRFVR**Stop**YG**St**
**op**ISSLVLRN**Stop**FL**Stop**INGCHRQNETTRYCDFSLSEDGIAEDSI**Stop**YLQLQ
YSDEVFLQLS**Stop**IVLCFVYIWQDHQTWFSSRCCYV**Stop**HPNPRVMetSGR**Stop**
DF**Stop**GRSFFYYMetVETGCPANVVTFTTLMetNGLCREGRVLQALALVDRMetVE
EGHQPDAVTYGTIVNGMetCKLGDTVSALNMetLRKMetDESQIKANVVIYSAIVDR
LCKDGNHIKAQNIFTEMetHEKGIFPNVLTYNCMetIDGYCSYGKWSDAEQLLRDMe
tIERNIDPDVVTFSALINAFVKEGKVSGAEELYREMetLRRNIFPTTITYSSMetID
GFCKHSRLEDAKHMetFDLMetVSKGCSPDIITLNTLIDGCCRAKRVDDGMetKLLH
EMetSRRGLVPDTVSYSILFTGSVKWGMetLMetLLKTFSRR**Stop**FLMetVCPLIS
**Stop**LVTLCWPVSARMetGS**Stop**KRRWKCLRFSRKVRWILILLLVTSSSMetECAR
VIRWTKHGICSIVSPSMetVWKLMetS**Stop**LTIY**Stop**SAYLSKKGTF**Stop**GLKI
FTWKCSVKV**Stop**FPVLSHITQW**Stop**MetGSANRTA**Stop**KRPDRWSIRWLVKAAP
LT**StopStop**PLVHSLKAIVRQEGLMetTD**Stop**SFSARCVKGD**Stop**LLTQLHTTL
**Stop**FMetGFVKWVILMetALKIYSRRWFLVVCVLIPLLSVVCWLVYVLRRNYKRD**S**
**top**QCWRICRKVWYVLYSLLYFRVLEMetNVYIYRALAGELLV**Stop**HPEKFATECL
SQI**Stop**DRKFYLNPL**Stop**IAWCVLLCRIMetNWRMetMetSERIGRFNAISLS**St**
**op**FHFYYN**Stop**K**Stop**MetIMetVLT**Stop**LFILDIYVFMet**Stop**WKKFQFSSSV
LSSKKYRFLRVVP

**Supplemental Figure 9. Translated protein sequence of pseudo-*PPR* with predicted ORFs (open reading frames).**

The pseudo-PPR gene sequence was translated to amino acid sequence using Expasy web server at default parameters. The ORFs (red color) was predicted using the Expasy online tool.

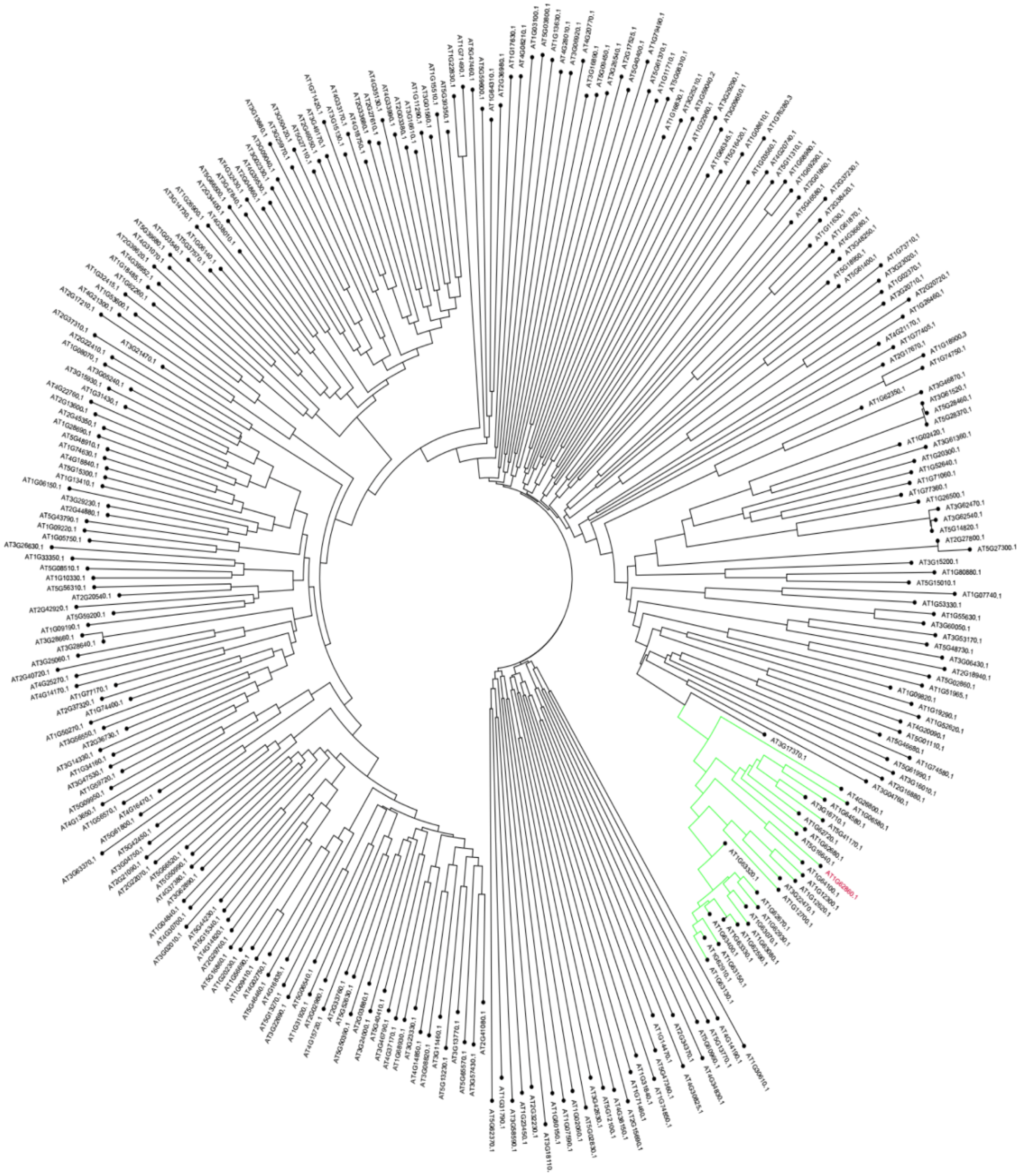

**Supplemental Figure 10. Phylogenetic analysis of *PPR* genes**

Unrooted Neighbor-Joining tree of 297 *PPR* gene sequences of *Arabidopsis thaliana* using MEGA6.0 at default parameter. The green branch indicates the PPRs with P-type motif forming a cluster with pseudo-*PPR* (red color).

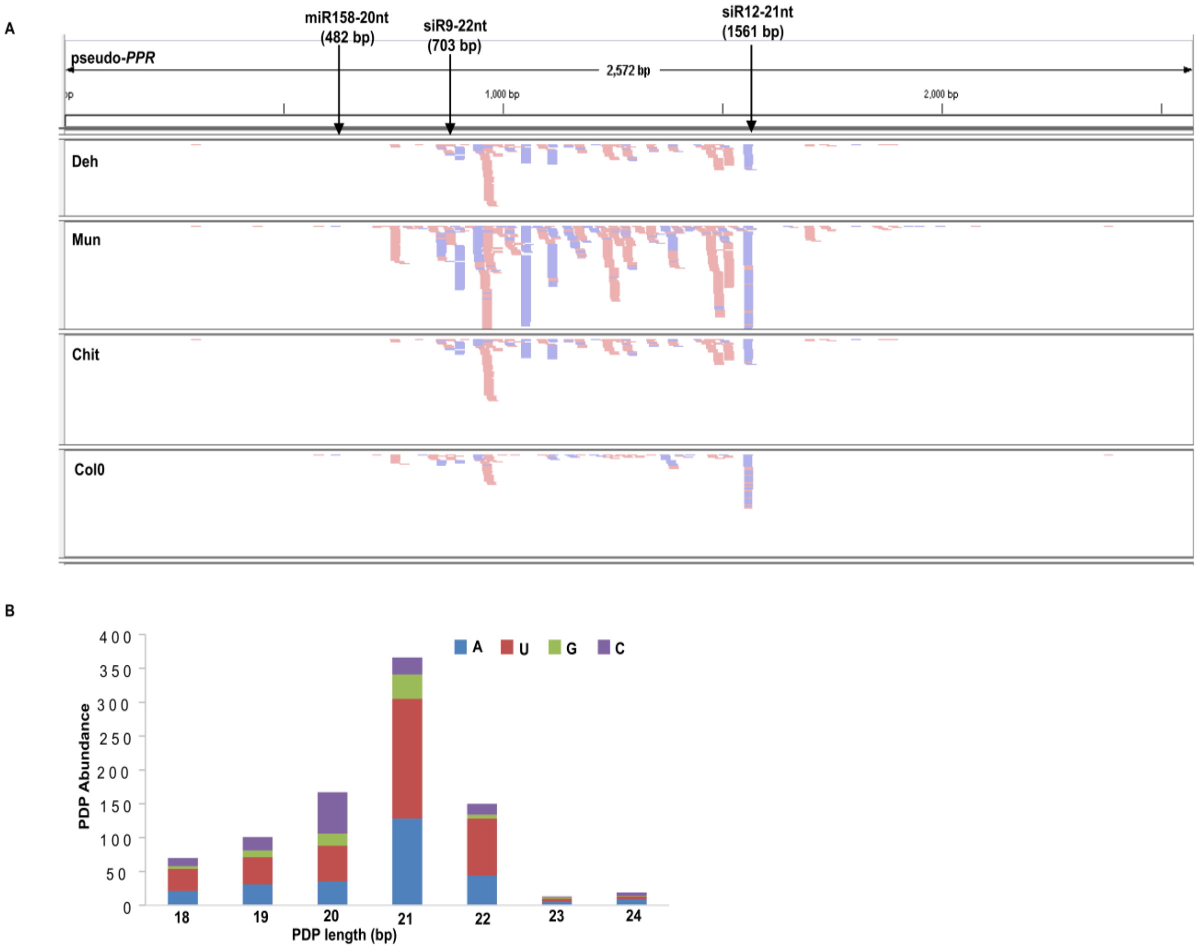


**Supplemental Figure 11. Abundance of phasiRNA (PDPs) in different populations**

- **(A)** The small RNA reads mapped onto pseudo-*PPR* from different populations (Deh, Mun, Chit and Col0) are shown in different tracks in Integrative Genomics Viewer (IGV). The cleavage site of miR158, siR9 and siR12 is marked by an arrow along with the nucleotide position. Red and blue blocks represent reads mapped in forward and reverse orientation, respectively.
- **(B)** Bar plot representing length distribution of the PDPs. The first nucleotide of the PDPs of different categories is depicted with different colors. 21 nucleotides long PDPs with 5'Uridine were predominant ones.

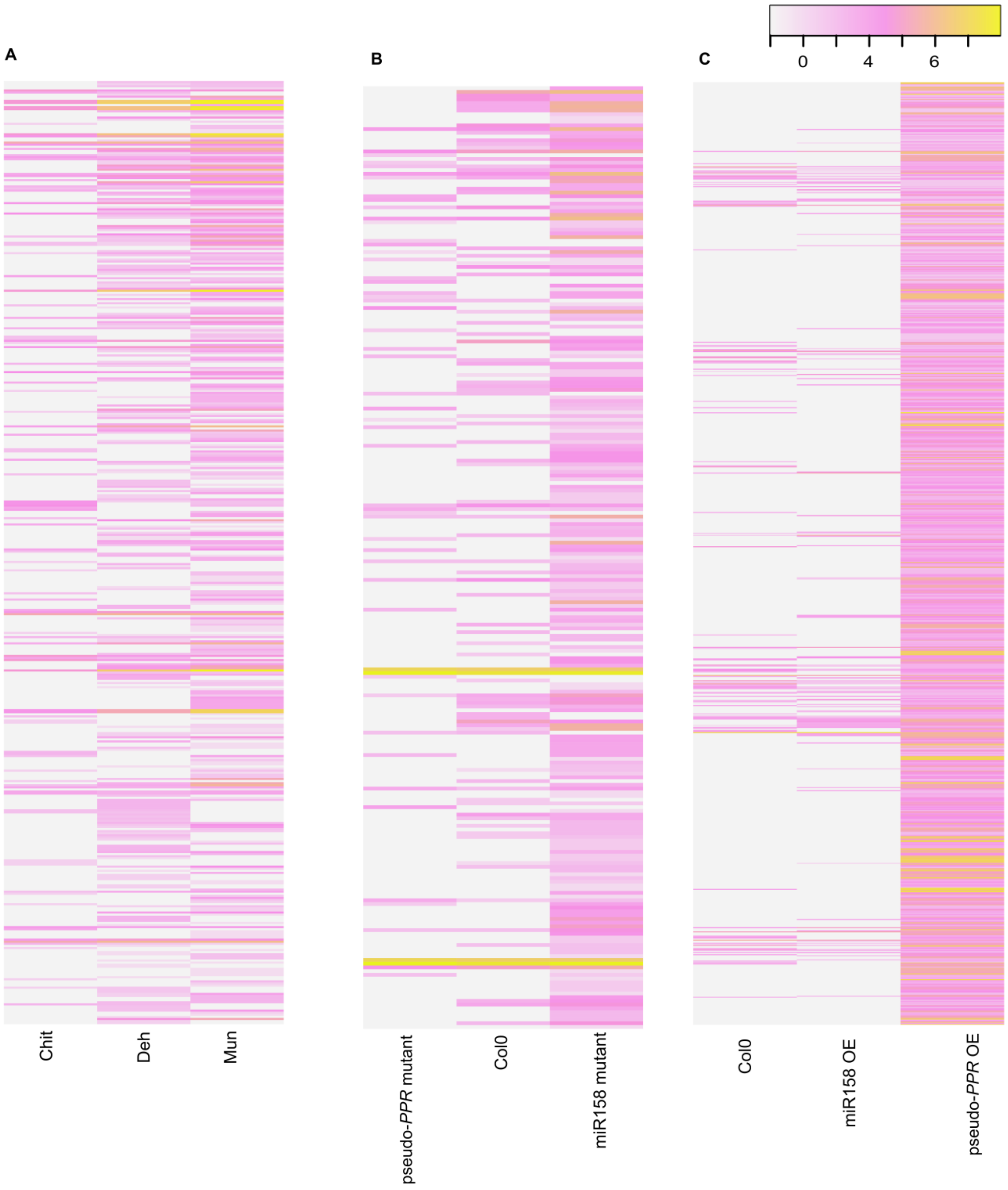


**Supplemental Figure 12. Differential expression of PDPs using log2 values of read counts from different plant samples**

Heat maps indicating differential expression of PDPs in **(A)** different west Himalayan populations (n=452) **(B)** in Col0, pseudo-*PPR* mutant and miR158 mutant **(C)** in Col0, miR158 OE and pseudo-*PPR* OE. The log2 values of read counts are represented by colour intensity scale.

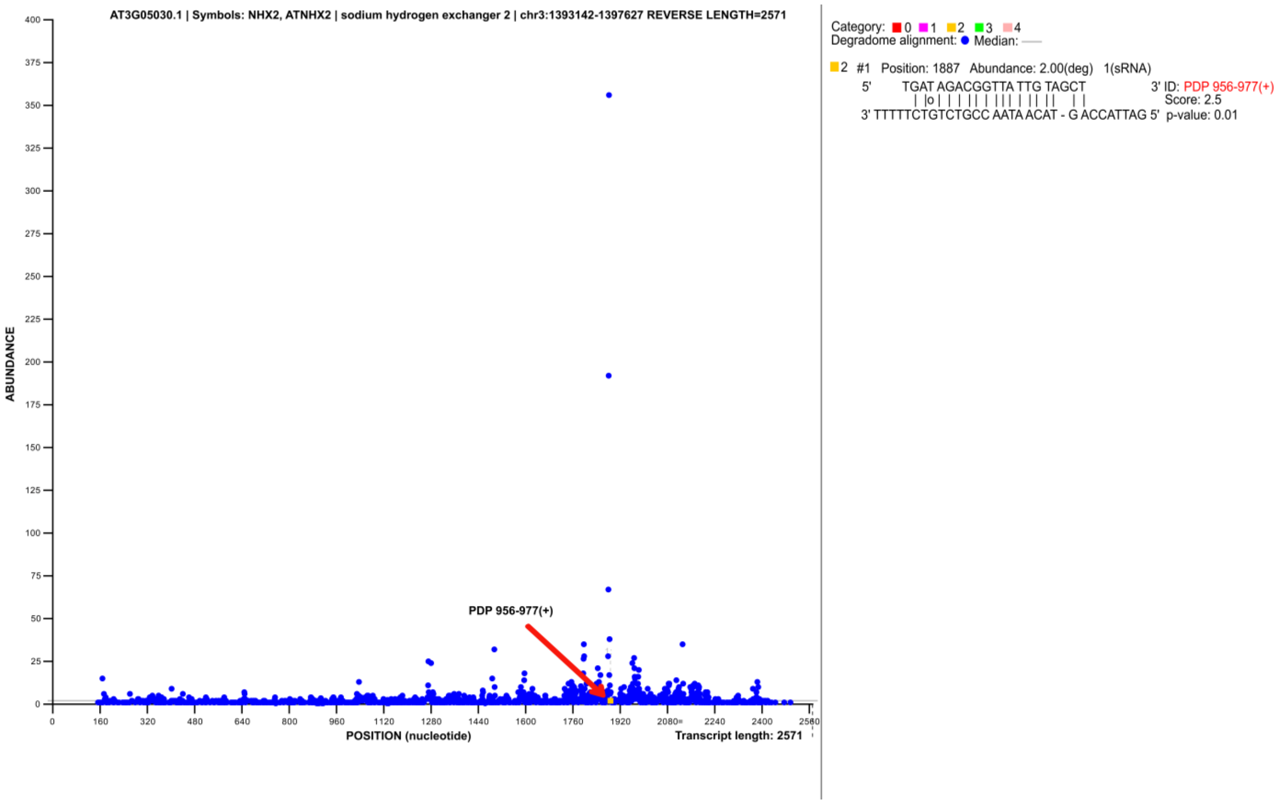


**Supplemental Figure 13. Target-plot (T-plot) of *NHX2***

T-plot of *NHX2* using degradome data. PDP 956-977(+) directed cleaved transcript (represented by an orange square) is marked by an arrow. The blue circles represent the fragments mapped on the pseudo-*PPR*. The X and Y axis represents nucleotide position on the targeted transcript and normalized read abundance (TPM, transcripts per million) of cleaved transcript detected in degradome-seq, respectively. mRNA:phasiRNA alignments along with p-value, alignment score and cleavage positions are also shown.

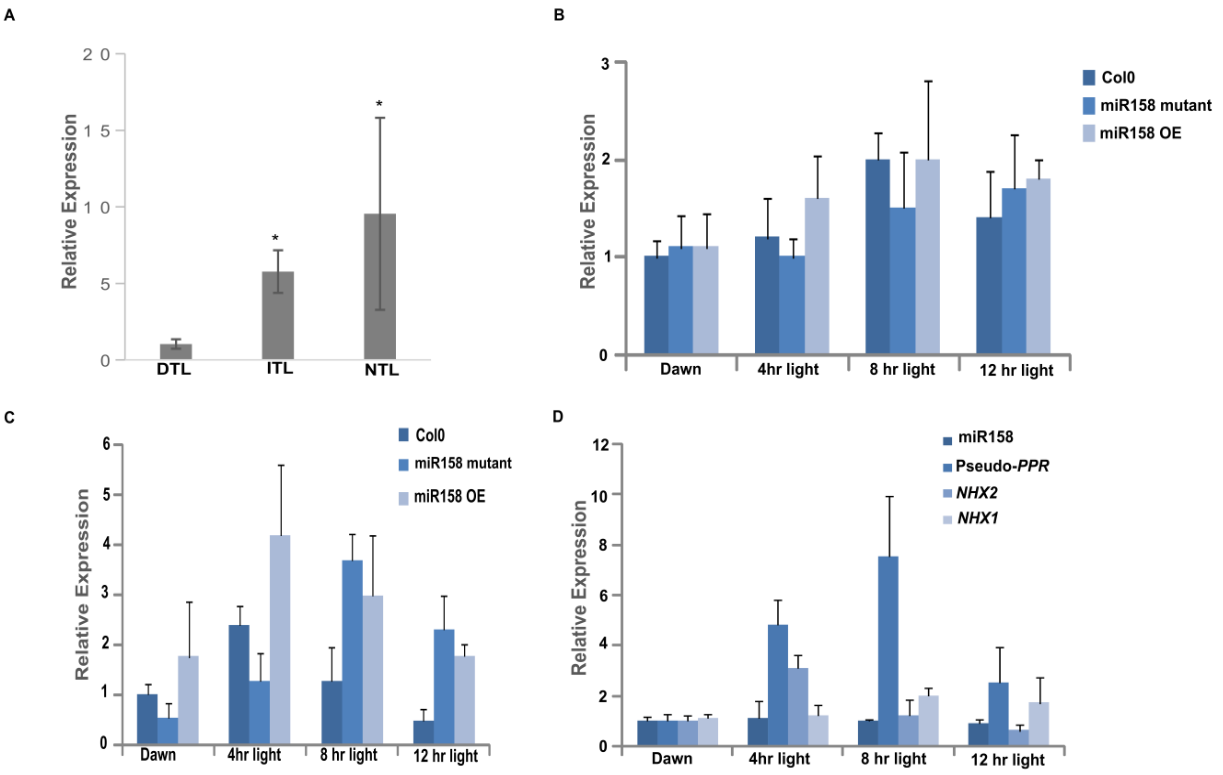


**Supplemental Figure 14. Relative expression pattern of *NHX2*, *NHX1*, pseudo-*PPR* and miR158 using qRT-PCR and stem-loop qRT-PCR**

**(A)** Expression pattern of *NHX2* in Indian west Himalayan *Arabidopsis* populations. The data represents the mean of three independent biological replicates and line above bars shows ± SD. * $P \leq 0.05$, (Student's t-test). Diurnal variation in relative expression of **(B)** *NHX1* and **(C)** *NHX2* in Col0, miR158 mutant and miR158 OE calculated using q-RT PCR. The data represent the mean of three biological replicates ± SD **(D)** Diurnal variation in relative expression pattern of miR158, pseudo-*PPR*, *NHX2* and *NHX1* in Col0 calculated using q-RT PCR. The data represent the mean of three biological replicates ± SD.

The expression data for miRNA and mRNA was normalized by the expression of 5s-rRNA and actin endogenous control, respectively and relative expression was calculated using $2^{-\Delta\Delta Ct}$ method.

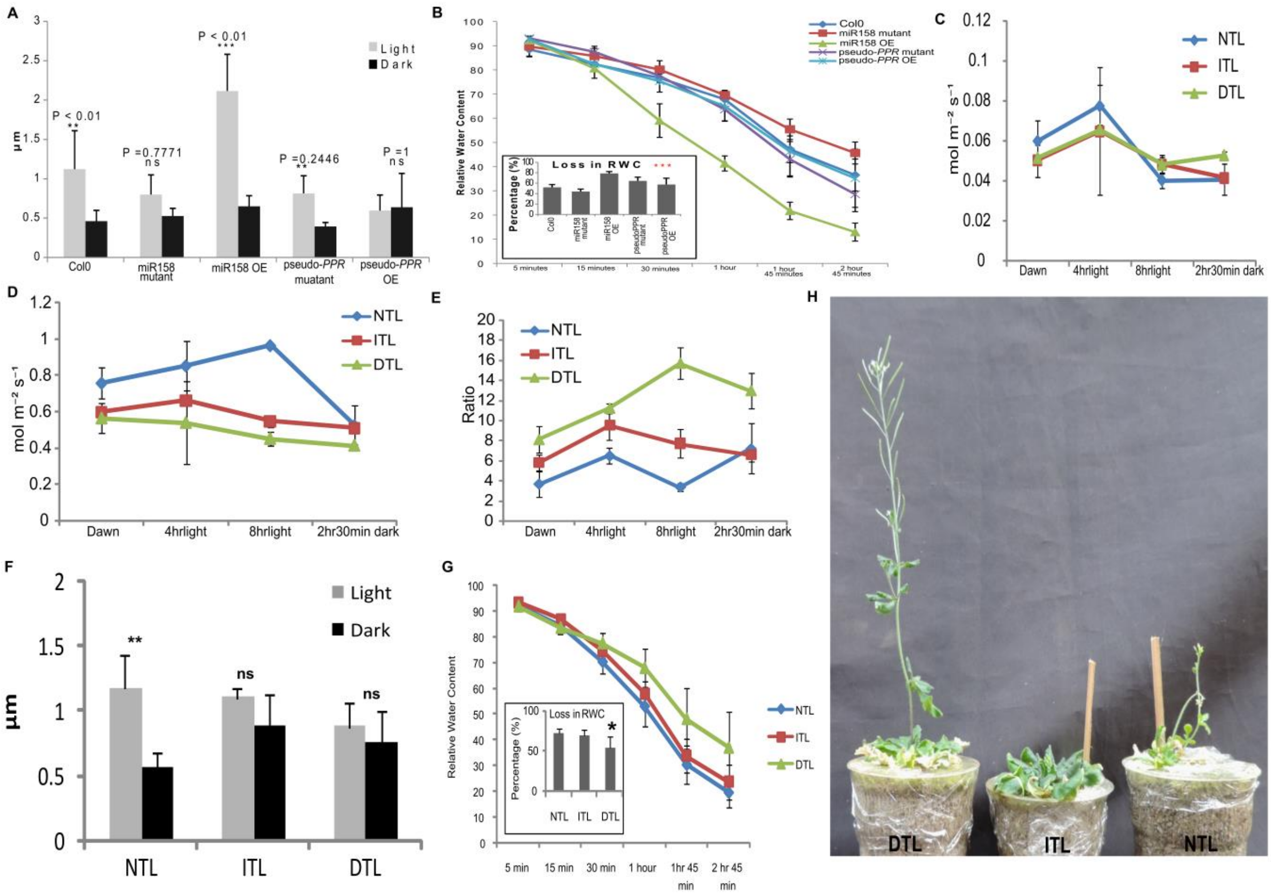


**Supplemental Figure 15. Physiological parameters of different plant samples**

- **(A)** Size of stomatal aperture of mutant and transgenic lines in Col-0 measured under light and 2 hr 30 min of dark conditions. The bar and the line above the bars represent the mean and ± SD, respectively. The values above the bars show P-value for significance using Students t-test.
- **(B)** Relative water content measured at different time points (inset bar plot representing total loss of water content, calculated as initial RWC (5min) - final RWC (2hr 45min)). Asterisk above the graph shows significance of variation using ANOVA (‘***’ P<0.001).

  Physiological parameters of NTL, ITL and DTL plants measured at different time points: **(C)** Stomatal conductance; **(D)** Transpiration rate **(E)** Water Use Efficiency. The data represents the mean of five biological replicated ± standard deviation.
- **(F)** Size of stomatal aperture of NTL, ITL and DTL measured under light and 2 hr 30 min of dark conditions. The bar and the line above the bars represent the mean and ± SD, respectively. The values above the bars show P-value for significance using Students t-test.
- **(G)** Relative water content measured at different time points (inset bar plot representing total loss of water content, calculated as initial RWC (5min) - final RWC (2hr 45min)). Asterisk above the graph shows significance of variation using ANOVA (‘***’ P<0.001).
- **(H)** Image showing DTL, ITL and NTL plants after 35 days of water holding.

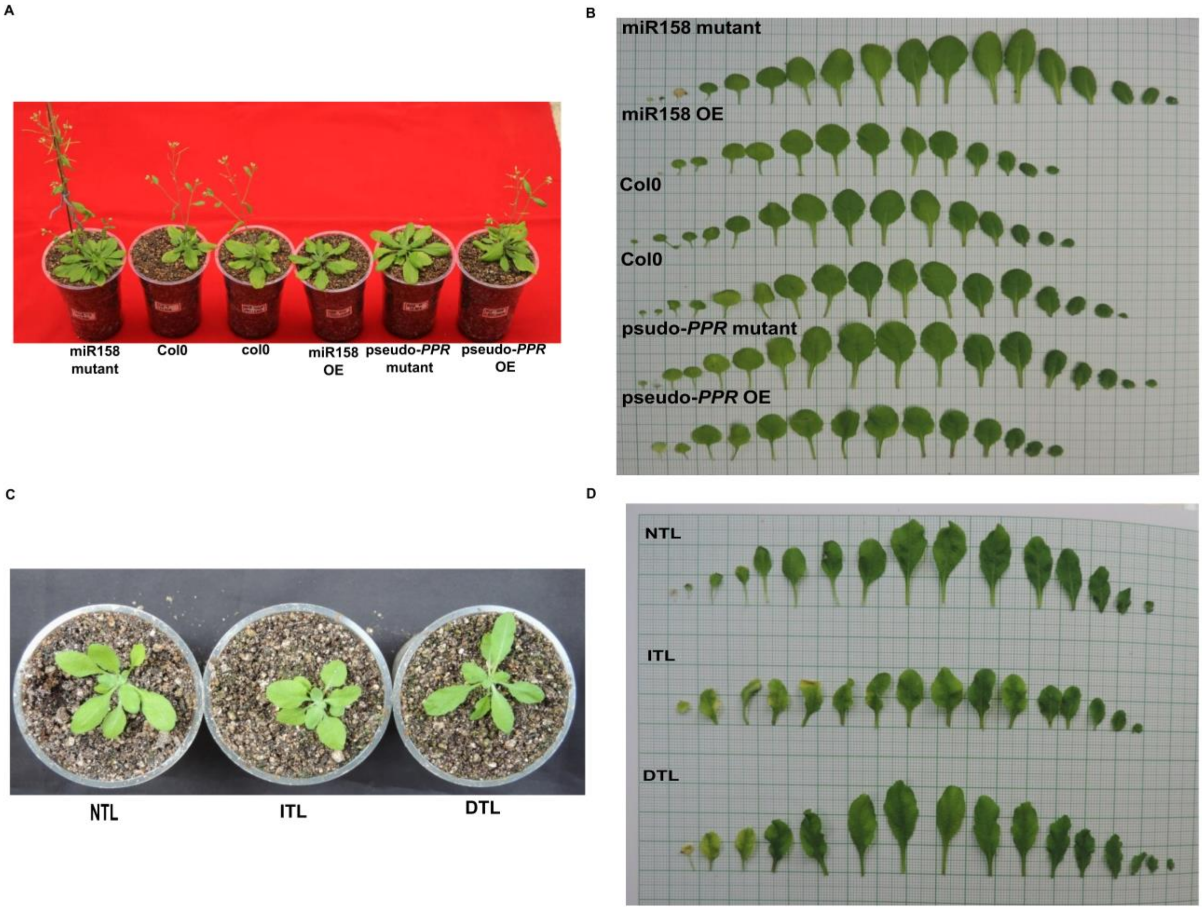


**Supplemental Figure 16. Growth and leaf size of different transgenics and Himalayan populations**

**(A)** Images showing miR158 and pseudo-*PPR* mutant with different transgenic lines in Col0 background. The image was taken 45 days post germination. **(B)** Serially arranged leaves of mutants and transgenic lines in Col0 background to depict variations in leaf morphology. The image was taken 30 days post germination. **(C)** Image of NTL, ITL and DTL plants at 30 days post germination. **(D)** Serially arranged leaves of NTL, ITL and DTL plants to depict variations in leaf morphology. The image was taken 30 days post germination.

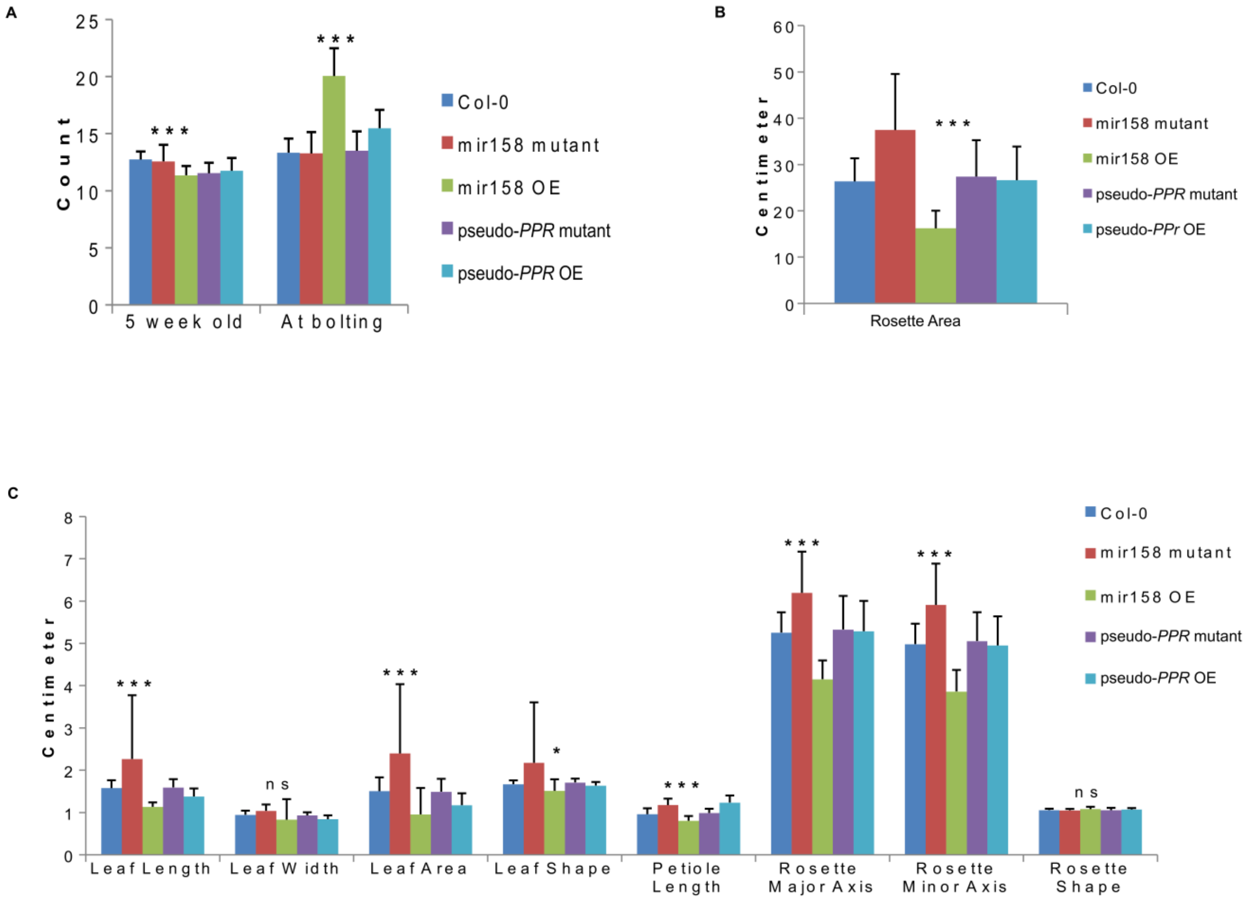


**Supplemental Figure 17. Morphological variations in different transgenic lines and Col0**

Bar plots representing morphological variations in mutant and transgenic lines in Col0 background. **(A)** Leaf count at two stages **(B)** Rosette area **(C)** Other leaf traits. The plot represents mean trait value with ±SD shown as line above the bars. The significance value is shown above the bars. ('***' P<0.001, '** P<0.01, '*' P<0.05, ns=not significant).

A CTCTTGTTGATCGGATGGTAGAA 711 **pseudo-*PPR***

|||||||||||||||||||||||

CTCTTGTTGATCGGATGGTAGAA 605 **TAS-2A**

B >AT1G62860.1 | Symbols: No Symbol | pseudogene of Pentatricopeptide repeat (PPR) superfamily protein

GCTCCTCCATTTGTTGTTGTTAGAGTTTAGAGACAACGAAAATCAGGCGAGTCGAAAATGTTGGCTAGGGTTTGCAG
ATCTGAATCATCTTCTGGTAATGCTGCTGCTGTTTCTGCGAGATTGTTCAGTACGAGATTAGTTCATCGCCGCGTCG
CCAAGAAAACTAGTAAGGTTGAAAAGAGTATGCCTTCTCTGAGGAGCTTTTGAAGGAGAGAGTTTGAAATTGAGAAG
TGGATCTCATTATATCAGAAGCTTAGATGATGCGATCGATTTGTTCGATGATATGGTTAGATCTCGTCCCTTGTACT
CAGAAATTGATTTCTGTAAATTAATGGGTGTCATCGTCAGAATGAAACAACCCGATATTGCGATTTCTCTCTATCAG
AAGATGGAATTGCGGAGGATTCCATTTGATATCTACAGCTTCAATATTCTGATGAAGTGTTTCTGCAGCTGTCATAA
ATTGTCCTTTGCTTTGTCTACATTTGGCAAGATCACCAAACTTGGTTTTCATCCCGATGTTGTTACGTTTAGCACCC
TAATCCACGGGTTATGTCTGGAAGGTAGGATTTCTGAGGCCGTAGCTTTTTTTATTATATGGTTGAAACGGGATGTC
CTGCAAATGTCGTAACGTTTACCACACTGATGAACGGTCTTTGCCGCGAGGGTCGAGTTCTCCAAGCGCTAG***CTNNN***
***NNNNNNNNGATGGTAGAA***GAAGGTCATCAACCTGACGCGGTTACTTACGGAACAATTGTCAATGGAATGTGTAAGTT
GGGTGACACTGTCTCGGCTTTGAATATGCTTAGGAAGATGGATGAAAGCCAAATCAAAGCCAATGTTGTAATCTATT
CTGCCATCGTTGATCGCCTTTGCAAGGACGGGAACCACATCAAGGCTCAAAATATTTTCACTGAAATGCATGAGAAA
GGTATTTTTCCCAATGTTCTCACTTACAATTGTATGATAGACGGTTATTGTAGCTATGGTAAATGGAGTGATGCGGA
GCAATTGCTGCGTGATATGATCGAAAGGAATATCGACCCTGATGTTGTTACTTTCAGCGCATTGATCAATGCATTTG
TCAAAGAAGGAAAGGTCTCTGGGGCTGAAGAATTATACCGTGAGATGCTTCGAAGGAATATATTTCCTACCACAATC
ACGTATAGTTCAATGATCGATGGATTTTGCAAGCATAGCCGTCTAGAAGACGCAAAGCACATGTTTGATTTGATGGT
TAGCAAGGGCTGCTCTCCGGATATAATTACTTTAAATACTCTCATAGATGGATGCTGTAGGGCGAAGAGGGTAGATG
ATGGAATGAAGCTTCTCCATGAGATGTCGAGAAGAGGATTAGTTCCTGATACCGTTTCTTACAGCATCTTATTCACG
GGTTCTGTCAAGTGGGGAATGTTAATGTTGCTCAAGACCTTTTCCAGGAGATGATTTCTAATGGTGTGTCCCCTGAT
ATCGTAACTTGTAACACTCTGCTGGCCGGTCTCTGCGAGAATGGGAAGTTAGAAAAGGCGTTGGAAATGTTTAAGGT
TTTCCAGAAAAGTAAGATGGATCTTGATACTGCTACTTGTAACATCATCATCAATGGAATGTGCAAGGGTAATAAGG
TGGACGAAGCATGGGATTTGTTCAATAGTCTCCCCGTCAATGGTGTGGAAACTGATGTCGTAACTTACAATATATTG
ATCGGCGTATTTGTCAAAGAAGGGAACTTTTTAAGGGCTGAAGATATTTACTTGGAAATGCTCTGTAAAGGTATAAT
TCCCAGTACTGTCACATATAACTCAATGGTAGATGGGTTCTGCAAACAGAACCGCCTAGAAGAGGCCAGACAGATGG
TCGATTCGATGGTTAGTGAAGGCTGCTCCCCTGACGTAGTGACCTTTAGTACACTCATTAAAGGCTATTGTAAGGCA
GGAAGGGTTGATGACGGATTAGAGCTTTTCAGCGAGATGTGTCAAAGGGGATTAGTTGCTGACACAATTACATACAA
CGCTTTGATTCATGGGTTTTGTAAAGTGGGTGATCTTAATGGCGCTCAAGATATATTCGAGGAGATGGTTTCTAGTG
GTGTGTGTCCTGATACCATTACTTTCCGTAGTATGCTGGCTGGTCTATGTACTAAGGCGGAACTACAAAAGGGATTG
ACAATGTTGGAGGATTTGCAGAAAAGTGTGGTATGTCCTCTACTCACTTTTGTACTTTCGAGTTTTGGAGATGAATG
TATATATATACAGAGCTTTGGCTGGAGAGCTTCTAGTGTAGCATCCGGAGAAGTTTGCAACTGAATGTCTAAGTCAA
ATTTAAGATAGAAAATTCTATCTTAACCCTCTCTAAATTGCTTGGTGTGTGTTGCTTTGCAGGATCATGAATTGGAG
GATGATGAGTGAAAGAATTGGAAGATTCAATGCCATTTCCCTTTCGTAATTCCATTTCTATTACAATTGAAAATGAA
TGATTATGGTTCTAACTTGATTATTCATTCTGGATATTTATGTGTTCATGTAGTGGAAAAAGTTTCAATTTTCAAGC
TCAGTTCTATCCTCAAAAAAGTATAGATTTCTGAGAGTTGTCCCGAG

**Supplemental Figure 18. Masked sequence of pseudo-*PPR* used for identifying pseudo-*PPR* derived phasiRNAs (PDPs)**

**(A)** 23 bp complementarity between pseudo-*PPR* and TAS-2 gene.

**(B)** Masked reference sequence of pseudo-*PPR* used for identifying PDPs. The 23 bp complementary region between TAS-2 and pseudo-*PPR* was masked by replacing the nucleotides lying in complementary region with N (shown as bold and underlined).

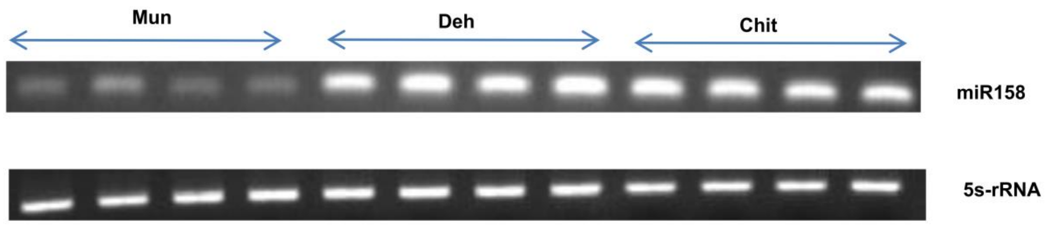


**Supplemental Figure 19. Gel Image of Endpoint PCR in the three populations**

Gel image showing the results for endpoint PCR of miR158 and endogenous control (5s-rRNA) used to check the specificity of the designed stem-loop primer of miR158. Similar amount of amplified products were loaded on 2% agarose gel. Presence of single band in each PCR reaction confirms the specificity of the primers as well as the variation in the expression pattern of miR158.

**Supplementary Table 1. Length distribution of expressed form of siR9 and siR12 in NTL, ITL and DTL.**

| | NTL | | DTL | | ITL | |
|---|---|---|---|---|---|---|
| **Length (nt)** | **siR9** | **siR12** | **siR9** | **siR12** | **siR9** | **siR12** |
| **18** | 50 | 18 | 102 | 29 | 53 | 5 |
| **19** | 481 | 61 | 680 | 66 | 682 | 20 |
| **20** | 2019 | 143 | 2076 | 164 | 2379 | 49 |
| **21** | 2258 | 3191 | 1843 | 2694 | 1215 | 1543 |
| **22** | 4235 | 232 | 3393 | 211 | 2273 | 149 |
| **23** | 325 | 27 | 266 | 34 | 270 | 18 |
| **24** | 137 | 8 | 100 | 9 | 123 | 6 |

22nt length is the most dominant form for siR9 which is highlighted in grey.
